# Maternally inherited siRNAs initiate piRNA cluster formation

**DOI:** 10.1101/2022.02.08.479612

**Authors:** Yicheng Luo, Peng He, Nivedita Kanrar, Katalin Fejes Toth, Alexei A. Aravin

**Affiliations:** Division of Biology and Biological Engineering, California Institute of Technology, Pasadena, CA 91125, USA; European Molecular Biology Laboratory, European Bioinformatics Institute (EMBL-EBI), Wellcome Genome Campus, Cambridge, UK

## Abstract

PIWI-interacting RNAs (piRNAs) guide repression of transposable elements in germlines of animals. In *Drosophila*, piRNAs are produced from heterochromatic genomic loci, called piRNA clusters, that act as a repositories of information about genome invaders. piRNA generation by dual-strand clusters depend on the chromatin-bound Rhino-Deadlock-Cutoff (RDC) complex, which is deposited on clusters guided by piRNAs, forming a feed-forward loop in which piRNAs promote their own biogenesis. However, how piRNA clusters are formed initially, before cognate piRNAs are present, remained unknown. Here we report spontaneous *de novo* formation of a piRNA cluster from repetitive transgenic sequences. We show that cluster formation occurs gradually over several generations and requires continuous trans-generational transmission of small RNAs from mothers to their progeny. We discovered that maternally-supplied siRNAs are responsible for triggering *de novo* cluster activation in progeny. In contrast, the siRNA pathway is dispensable for cluster function after its establishment. These results revealed an unexpected cross-talk between the siRNA and piRNA pathways and suggest a mechanism for *de novo* formation of piRNA clusters triggered by production of siRNAs.

**Highlights:**

- A transcribed repetitive transgene is spontaneously converted into dual-strand piRNA cluster
- Establishment of piRNA cluster occurs over multiple generations and requires cytoplasmic inheritance of cognate small RNA from mothers
- Cognate siRNAs initiate the activation of piRNA cluster, but are dispensable after its establishment

## Introduction

Binary complexes of small non-coding RNAs and Argonaute (Ago) proteins play essential roles in regulating gene expression and suppressing foreign and selfish nucleic acids. Small RNAs guide the Argonautes to complementary RNA targets upon which Agos either cleave the target or recruit additional factors to repress them by other mechanisms. Despite the common architecture of Ago-small RNA complexes, there are three distinct classes of small RNAs - siRNA, miRNA and piRNA - that differ in their biogenesis, functions as well as the specific members of the Ago family they partner with.

Both siRNA and piRNA were reported to suppress activity of endogenous (transposable elements and other selfish genes) and exogenous (viruses) elements in various animal species (Aravin et al., 2006; Brennecke et al., 2007; Gammon and Mello, 2015; Gitlin et al., 2002; Lindbo et al., 1993; Vance and Vaucheret, 2001; Voinnet et al., 1999), however, the two pathways are believed to work independently of each other. Despite the complete abrogation of the siRNA pathway in Ago2 deficient flies, these flies are viable and fertile and show only mild activation of a few transposons in somatic tissues (Czech et al., 2008; Ghildiyal et al., 2008; Kawamura et al., 2008; Pelisson et al., 2007). In contrast, flies deficient for piRNA pathway components demonstrate strong activation of multiple TE families in the germline associated with DNA damage and complete sterility (Brennecke et al., 2007; Sienski et al., 2015; Vagin et al., 2006; Yu et al., 2015b).

siRNAs are processed from double-stranded or hairpin precursors by the Dicer nuclease and then loaded into their Ago protein partner (Bernstein et al., 2001; Matranga et al., 2005; Rand et al., 2005). siRNA/Ago complexes cleave complementary RNA targets in the cytoplasm leading to their degradation (Matranga et al., 2005; Rand et al., 2005). In *Drosophila*, siRNAs associate exclusively with Ago2 and mutation of Ago2 abrogates siRNA-guided repression (Czech et al., 2008; Ghildiyal et al., 2008; Kawamura et al., 2008; Okamura et al., 2004; Vagin et al., 2006). The biogenesis and function of piRNAs, however, is much more complex than that of siRNAs. piRNA processing is independent of Dicer but involves multiple other proteins (Andersen et al., 2017; Chen et al., 2016; ElMaghraby et al., 2019; Mohn et al., 2014; Vagin et al., 2006; Zhang et al., 2014). piRNAs are expressed in the germline and closely associated somatic cells of the ovary and testis and are loaded into a distinct clade of Argonautes called Piwi proteins, which in flies consist of Piwi, Aub and Ago3 (Brennecke et al., 2007; Saito et al., 2006; Vagin et al., 2006). Similar to siRNAs, cytoplasmic Piwi/piRNA complexes cleave complementary RNA targets, however, in addition to target degradation, this process also amplifies piRNAs through the so-called ping-pong cycle (Aravin et al., 2008; Brennecke et al., 2007; Gunawardane et al., 2007), thereby connecting their function to their own biogenesis. Furthermore, both flies and mice harbor a nuclear Piwi/piRNA complex capable of transcriptional repression through recognition of nascent transcripts followed by recruitment of a chromatin modifying machinery (Aravin et al., 2008; Le Thomas et al., 2013; Ninova et al., 2020; Sienski et al., 2015; Sienski et al., 2012; Yu et al., 2015b).

siRNA precursors are recognized by their double-stranded nature. Similar to siRNAs, piRNAs are also processed from longer RNA precursors, however these transcripts are single-stranded and lack distinct sequence and structure motifs, raising the question of how they are recognized and channeled into the processing machinery. In *Drosophila*, the chromatin-bound Rhino-Deadlock-Cutoff (RDC) protein complex marks dual-strand piRNA clusters, genomic regions that generate the majority of piRNAs in the germline (Klattenhoff et al., 2009; Zhang et al., 2014; Le Thomas et al., 2014b; Mohn et al., 2014; Chen et al., 2016). RDC is required for transcription of piRNA precursors by promoting initiation (Andersen et al., 2017; Mohn et al., 2014) and suppressing premature termination (Chen et al., 2016). RDC also promotes loading of the RNA-binding TREX (Hur et al., 2016) and Nxf3-Nxt1 (ElMaghraby et al., 2019) RNA export complexes on pre-piRNA. Loading of Nxf3-Nxt1 was proposed to channel precursors to the cytoplasmic processing machinery (ElMaghraby et al., 2019). Thus, RDC binding to chromatin of piRNA clusters might be both necessary and sufficient to sustain piRNA biogenesis.

Though the process of RDC deposition on chromatin is not completely understood, it seems to be guided by piRNAs, as the nuclear Piwi/piRNA complex directs deposition of the H3K9me2/3 mark, which in turn provides an anchor for RDC binding (Akkouche et al., 2017; Mohn et al., 2014; Sienski et al., 2015; Yu et al., 2015b). Several studies demonstrate the critical role of cytoplasmic piRNA inheritance from the mother to the progeny in initiating piRNA production (Casier et al., 2019; de Vanssay et al., 2012; Le Thomas et al., 2014b). Together these findings suggest that piRNA biogenesis is governed by a trans-generational feed-forward loop in which piRNA biogenesis is promoted by RDC complex, which in turn is deposited on chromatin guided by cytoplasmically inherited piRNAs. This feed-forward loop explains how piRNA profiles are maintained through generations. However, in order to adapt to new transposon invasions, the pathway must be able to generate novel piRNAs. How novel piRNA clusters arise is poorly understood.

Here we describe the *de novo* formation of a piRNA cluster over several generations. This process is accompanied by increasing piRNA levels and accumulation of the H3K9me3 mark and Rhi on cluster chromatin and requires continuous, maternal trans-generational cytoplasmic transmission of small RNAs. We found that cognate siRNAs trigger initial cluster activation, however, siRNA are dispensable after the cluster is established. Our results point to a tight cooperation between the siRNA and piRNA pathways in the fight against genome invaders and suggest that transposons are first detected by the siRNA pathway, which activates a robust piRNA response.

## Results

### Reporters inserted in the dual-strand 42AB cluster are repressed by piRNA, while reporters in the uni-strand 20A cluster disrupt cluster expression

To understand how new insertions into piRNA clusters are regulated, we integrated reporters into the two types of piRNA clusters – uni-strand and dual-strand clusters. We employed recombinase-mediated cassette exchange (RMCE) to integrate a reporter into specific genomic sites using a collection of *Minos*-mediated integration cassette (MiMIC) containing *D. melanogaster* stocks. The reporter encodes nuclear EGFP expressed under control of the ubiquitin (*ubi-p63E*) gene promoter, which drives expression in both somatic and germ cells (Fig. 1A). Using RMCE we inserted reporters into the major dual-strand cluster 42AB and the uni-strand cluster 20A (Fig. 1B). In the 20A cluster, the reporter was integrated 2.5 kb downstream of the cluster promoter and two reporter orientations were obtained. Reporters inserted in both clusters were expressed in somatic follicular cells of the fly ovary (Fig.1C), as well as in other somatic tissues (data not shown), indicating that the transgenes are functional and that integration into the repeat-rich cluster environments is compatible with reporter expression in somatic cells. Flies with reporters in the uni-strand 20A cluster, inserted in either orientation, also expressed GFP in the germline. Unlike 20A cluster transcripts, which localized to the nucleus, GFP mRNA was predominantly in the cytoplasm. In contrast, although fluorescent in situ hybridization revealed that both strands of the 42AB reporter sequence were transcribed, GFP protein was not expressed from the 42AB cluster insertion in the germline. Similar to native 42AB cluster transcripts, RNA transcribed from both strands of the 42AB reporter was concentrated in the nucleus.

**Figure 1.**
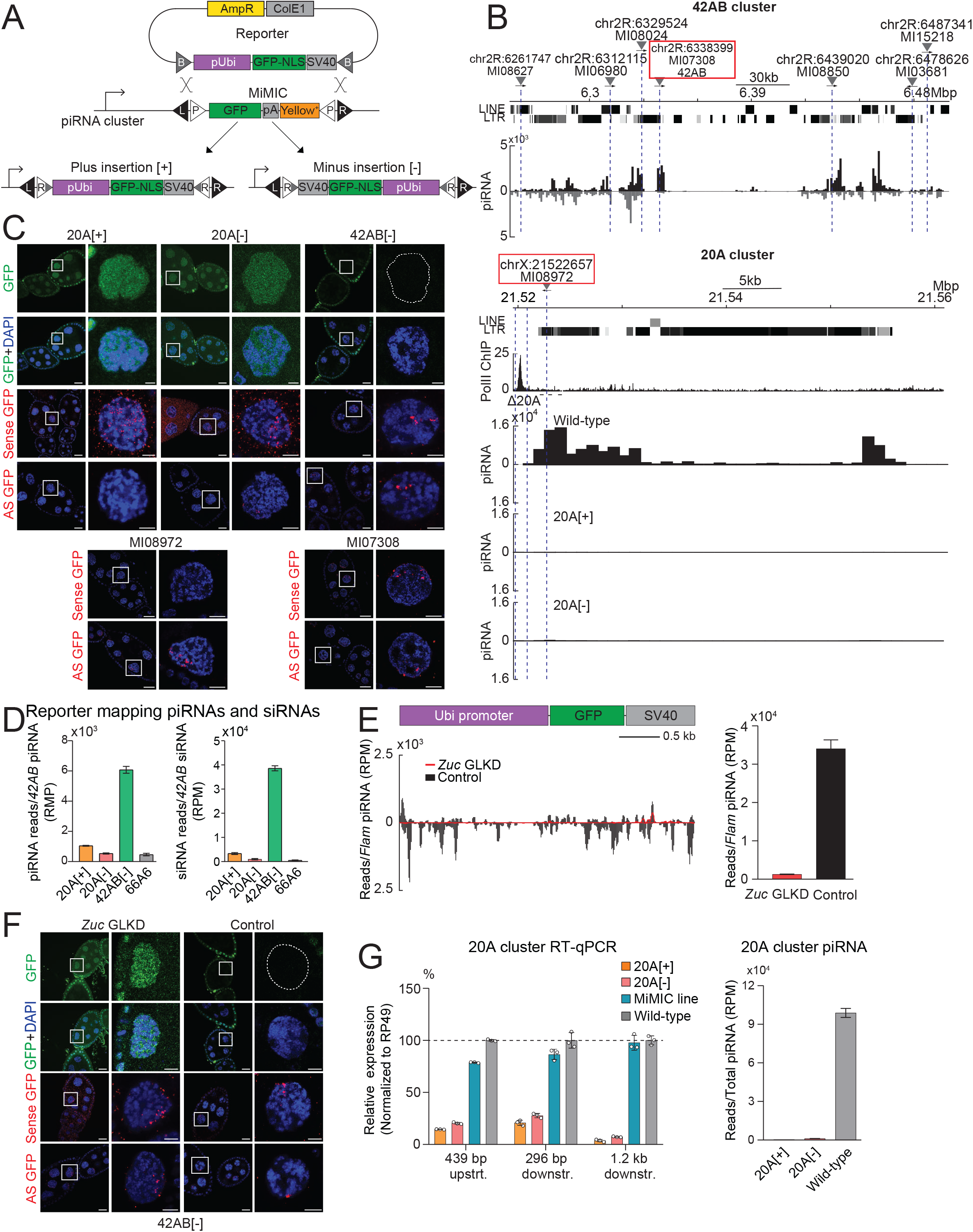
Reporters integrated in uni- and dual-strand clusters have different expression and effects on cluster activity. **(A) Scheme of reporter integration into piRNA clusters using recombinase-mediated cassette exchange (RMCE) to replace Minos-mediated integration cassettes (MiMIC)**. MiMICs contain GFP sequence but lack a promoter. In contrast, the reporter encodes nuclear EGFP expressed under control of the *Ubiquitin* gene promoter. RMCE can result in integration of the reporter in either orientation, which can be discriminated by genomic PCR. **(B) Profiles of the dual-strand cluster 42AB (top) and uni-strand cluster 20A (bottom)**. Shown are profiles of uniquely-mapping piRNAs, positions of the putative promoter (Pol II ChIP peak) of the 20A cluster and positions of the reporters. 20A cluster piRNA levels are dramatically reduced in ovaries of 20A[+] and 20A[-] flies. Number of piRNA reads mapped to the 20A cluster is normalized to total piRNA read count. **(C) Expression of reporters integrated in 20A and 42AB clusters**. Expression of GFP protein and sense and antisense RNA (by RNA FISH) in ovaries of flies with reporter insertions in the 20A and 42AB clusters. In all reporter lines GFP protein is expressed in somatic follicular cells, however, GFP is not expressed in germline (nurse) cells in the 42AB reporter line. Sense GFP RNA is produced in both 20A[+] and 20A[-] flies and it is predominantly localized in the cytoplasm of nurse cells. This cytoplasmic localization is different from the nuclear localization of exclusively antisense RNA produced by the promoterless MiMIC sequence integrated in the same site *(MI08972)*. The 42AB reporter generates transcripts from both strands and both sense and antisense RNA are localized in the nucleus, similar to transcripts produced by the promoterless MiMIC sequence integrated in the same site *(MI07308)*. Scale bar is 20µm and 2µm for egg chamber and single nurse cell nuclei, respectively. **(D) piRNAs and siRNA are generated from the reporter in the 42AB cluster, but not the reporter inserted into 20A or the control (non-cluster) locus**. Number of the small RNA reads mapping to the reporter were normalized to reads mapping to the 42AB cluster. Error bars indicate standard deviation of two biological replicates. **(E) Knockdown of *Zuc* eliminates 42AB reporter piRNA**. Shown are piRNA profiles along the reporter in ovaries of control flies (white GLKD, black) and upon GLKD of Zuc GLKD driven by nos-Gal4 (red). Bar graph shows number of piRNA reads mapped to the reporter normalized to *flamenco*-derived piRNAs, which are not affected by *Zuc* knockdown. Error bars indicate standard deviation of two biological replicates. **(F) Derepression of 42AB reporter expression upon *Zuc* GLKD**. Shown are GFP protein expression and FISH signal for both strands of the reporter. Note appearance of sense GFP transcripts in the cytoplasm upon *Zuc* GLKD. Scale bar is 20µm and 2µm for egg chambers and single nurse cell nuclei, respectively. **(G) Ovarian expression of the 20A cluster is suppressed upon reporter integration**. (Left) 20A cluster transcripts were measured by RT-qPCR using primers 439 bp upstream as well as 296 bp and 1217 bp downstream of the reporter insertion site. Cluster transcripts were normalized to *rp49* mRNA. Error bars indicate the standard deviation of three biological replicates. (Right) Expression of 20A cluster piRNA. piRNA reads uniquely-mapping to 20A cluster were normalized to total piRNA read count. Error bars indicate standard deviation of two biological replicates.

The exclusive repression of the 42AB reporter in the germline, where the 42AB cluster is active, suggest that repression might occur in a piRNA-dependent fashion. To explore if reporter sequences generate piRNAs, we cloned small RNA libraries from ovaries of transgenic animals. Analysis of the libraries revealed that reporter-derived piRNAs were abundant in flies with 42AB cluster insertions but not in flies with reporters in cluster 20A and in a control non-cluster 66A6 region (Fig. 1D). The 42AB reporter-derived piRNAs have the expected bias (69.95%) for U in the first position. Though piRNA were derived from both strands of the 42AB reporter sequence, 2.45-fold more piRNA are in antisense orientation relative to the GFP mRNA (Fig. 1E), indicating that they are not processed from reporter mRNA. In contrast, the few RNA reads derived from reporters inserted in 20A and the non-cluster region were predominantly in sense orientation and did not have a U-bias, indicating that they likely represent mRNA degradation products.

To explore whether the repression of GFP reporter inserted into the 42AB cluster depends on piRNAs, we knock-down Zucchini (Zuc), a critical piRNA biogenesis factor (Ipsaro et al., 2012; Nishimasu et al., 2012), in the germline. Depletion of Zuc led to strong (26.4-fold) reduction in the level of piRNAs mapping to the 42AB reporter (Fig. 1E) and to its derepression (Fig.1F). GFP protein expression upon Zuc GLKD correlated with the appearance of sense reporter transcripts in the cytoplasm of nurse cells, while antisense RNA remained in the nucleus. Thus, insertion of a gene into the 42AB cluster leads to generation of abundant piRNAs that trigger its repression.

To understand why 20A reporters do not produce piRNAs, we analyzed expression of 20A cluster transcripts by RT-qPCR using primer sets designed to detect cluster transcripts upstream and downstream of the reporter insertion site. Surprisingly, we found that the abundance of 20A cluster transcripts was strongly (> 20-fold) reduced in flies with reporter insertions compared to both wild-type flies and the original MIMIC flies used for RMCE (Fig. 1G). In agreement with the decreased cluster transcript level, 20A piRNA levels dropped 84- and 321-fold in 20A[-] and 20A[+] flies, respectively, throughout the whole cluster as far as 38 kb from the insertion site (Fig. 1B, G). In contrast, the insertion in the 42AB cluster did not affect piRNA level from this cluster (data not shown). As the original MIMIC line contains a promoterless insertion in the same site as the reporter lines, this result suggests that cluster expression is disrupted by reporter transcription rather than insertion of a heterologous sequence *per se*. Thus, insertion of an actively transcribed gene in the 20A cluster close to its promoter disrupts cluster expression.

### An unusual reporter behaves like a *bona fide* dual-strand piRNA cluster that is enriched in Rhi and depends on it to generate piRNA

Replacement of the MIMIC cassette with the reporter through recombinase-mediated cassette exchange (RMCE) leads to random orientation of the inserted sequence. Each replacement experiment generates several independent *Drosophila* lines that we maintained and in which we determined cassette orientation. Unexpectedly, we found that one of the lines with insertion into the 20A cluster has distinct properties. Unlike other lines that maintained reporter expression in both the soma and the germline, this line, which we dubbed 20A-X, lost germline expression after several months of propagation (Fig. 2A). The somatic GFP expression, which we also confirmed by detecting abundant cytoplasmic GFP mRNA in follicular cells by in situ hybridization, argues against genetic damage of the reporter cassette. GFP RNA was also detected in germline nurse cells, however, unlike 20A[+] and 20A[-] reporters, transcripts from 20A-X localized exclusively to the nuclei (Fig. 2A). Thus, unlike other 20A insertion lines and similar to insertions in 42AB, 20A-X shows normal GFP expression in follicular cells and strong GFP repression in the germline.

**Figure 2.**
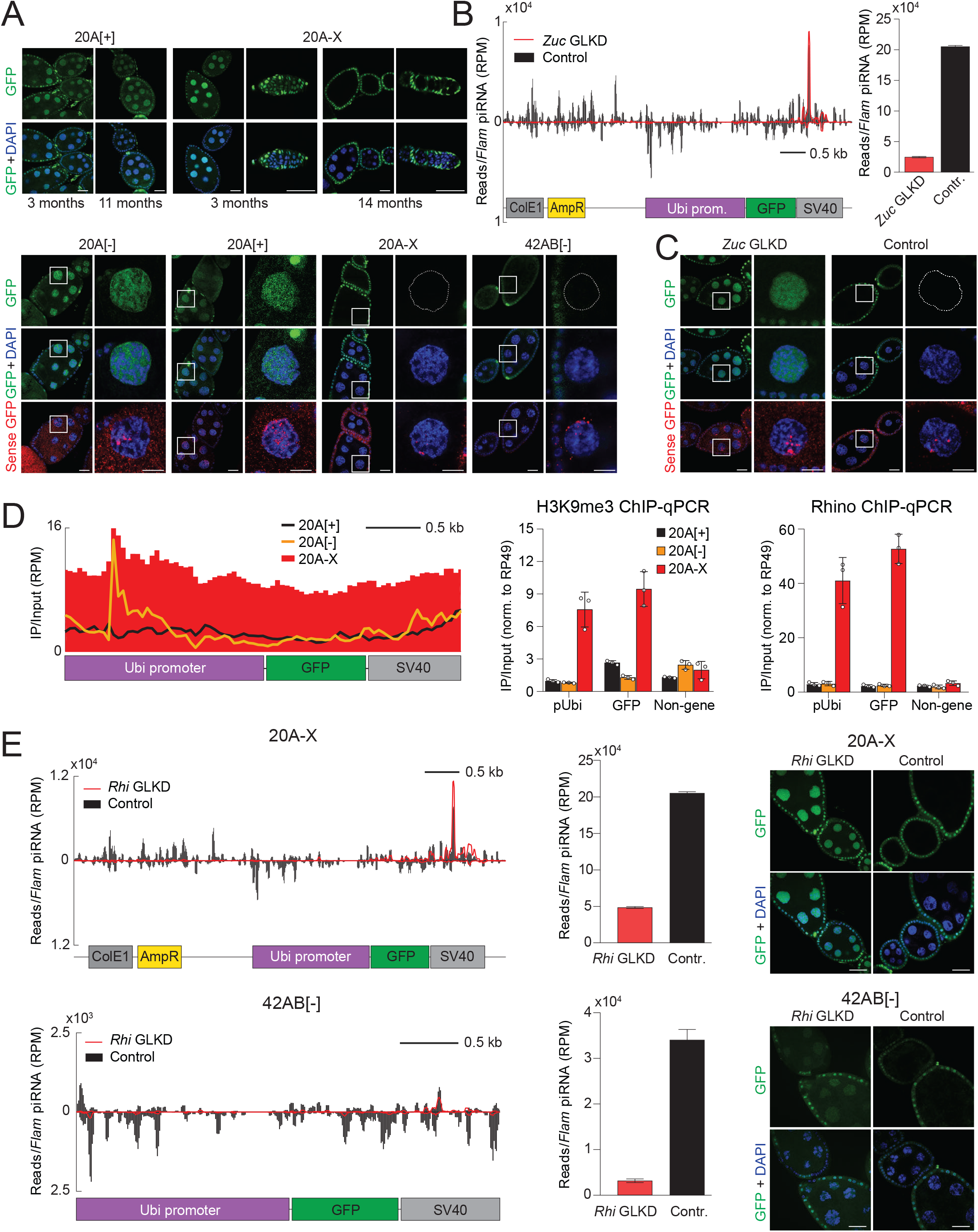
An unusual reporter insertion in the 20A cluster is a dual-stand piRNA cluster that generates piRNAs in a Rhi-dependent manner. **(A) The 20A-X reporter is silenced in the germline after several months of maintaining this line**. Top: GFP protein and sense RNA expression in ovaries of 20A[+] and 20A-X flies at different times after establishment of lines by RMCE. Bottom: GFP protein and sense RNA expression in ovaries of 20A[+], 20A[-], 20A-X and 42AB[-] flies 14 months (20A[+], 20A[-] and 20A-X) and 1 month (42AB[-]) after establishment of lines. Scale bar is 20µm and 2µm for egg chambers and single nurse cell nuclei, respectively. **(B) 20A-X generates piRNAs in a Zuc-dependent manner**. Shown are profiles of piRNAs mapping to the 20A-X reporter in control (*white* GLKD) and upon *Zuc* GLKD driven by nos-GAL4. 20A-X piRNAs were normalized to piRNA reads mapping to the *flam* cluster. Error bars indicate standard deviation of two biological replicates. **(C) *Zuc* knockdown releases 20A-X repression in the germline**. Shown are expression of GFP protein and sense RNA in control (*white* GLKD) and upon *Zuc* GLKD driven by nos-GAL4. Note that although GFP protein is not expressed in control flies, sense RNA is present in the nucleus. *Zuc* GLKD causes accumulation of RNA in the cytoplasm and leads to GFP protein expression. Scale bar is 20µm and 2µm for egg chambers and single nurse cell nuclei, respectively. **(D) 20A-X, but not other 20A reporters are enriched in Rhino and the H3K9me3 mark**. Left: Rhino ChIP-seq profiles (mean of two biological replicates) on 20A-X, 20A[+] (black) and 20A[-] (yellow) reporters. Right: Rhino and H3K9me3 enrichment on 20A reporter (Ubi and GFP region) as well as control locus (chr 2L: 968088-968187, dm6) were determined by ChIP-qPCR. Only the 20A-X reporter is enriched in Rhi and H3K9me3. ChIP signals were normalized to the *rp49* gene. Error bars indicate the standard deviation of three biological replicates. **(E) Rhino knockdown reduces 20A-X piRNA and releases its silencing in the germline**. Left: Shown are piRNA profiles over 20A-X and 42AB reporters in ovaries of control flies (*white* GLKD) and upon *Rhi* GLKD driven by nos-GAL4. Reporter piRNAs were normalized to piRNA reads mapping to the *flam* cluster. Error bars indicate standard deviation of two biological replicates. Right: GFP protein expression in 20A-X and 42AB ovaries upon *Rhi* GLKD. Scale bar is 20µm.

To test involvement of piRNAs in repression of the 20A-X reporter in the germline, we cloned and analyzed small RNA libraries from ovaries of 20A-X flies. This analysis revealed abundant piRNAs and siRNAs corresponding to the reporter sequence indicating that 20A-X is active as a piRNA producing locus. In fact, 20A-X generates 20.3-fold more piRNAs than the 42AB reporter. Germline knockdown of the piRNA biogenesis factor Zuc led to loss of 20A-X piRNAs (Fig. 2B). Zuc GLKD also led to release of the germline reporter repression as demonstrated by GFP protein expression and detection of sense RNA in the cytoplasm (Fig. 2C) indicating that, similar to 42AB insertions, repression of 20A-X is piRNA-dependent.

Several studies revealed essential differences between uni-strand and dual-strand piRNA clusters(Chen et al., 2016; Goriaux et al., 2014; Mohn et al., 2014). Dual-strand clusters, such as 42AB, are active exclusively in the germline and their transcription, nuclear processing and export require the Rhino-Deadlock-Cutoff (RDC) complex, which is anchored to these regions by the H3K9me3 histone mark(Le Thomas et al., 2014b; Mohn et al., 2014; Zhang et al., 2014). In contrast, uni-strand clusters, such as flamenco and 20A, do not depend on RDC and the H3K9me3 mark and can be active in the soma. To explore if 20A-X functions as a uni- or dual-strand piRNA cluster, we analyzed Rhino binding and H3K9me3 enrichment. ChIP-qPCR and ChIP-seq analyses revealed that, in contrast to the native 20A cluster and 20A[+] and 20A[-] reporters, 20A-X is strongly enriched in both Rhi and the H3K9me3 mark (Fig. 2D), suggesting that it acts as a dual-strand piRNA cluster.

To test whether 20A-X activity depends on RDC, we analyzed GFP repression and small RNA profile upon germline knockdown of Rhi. As expected, Rhi GLKD reduces the level of piRNAs generated from the 42AB reporter (Fig. 2E). Rhi GLKD also caused 4.2-fold reduction in 20A-X piRNA levels and released repression of GFP in germline of 20A-X flies (Fig. 2E). Taken together, our results indicate that 20A-X acts as a genuine dual-strand piRNA cluster that generates piRNAs in Rhi-dependent manner.

### The repetitive organization of the 20A-X locus correlates with its function as dual-strand piRNA cluster

We employed several approaches to understand how 20A-X differs from 20A[+] and 20A[-] reporters. Genomic PCR of flanking regions suggested that 20A-X harbors the reporter cassette in the correct site in the 20A cluster (Fig. 3A). In fact, similar to other 20A reporters, expression of 20A cluster transcripts is decreased in 20A-X flies (Fig. S3A). To confirm the insertion site, we employed *in situ* hybridization on salivary gland polytene chromosomes using the Ubi-GFP reporter sequence as a probe. *In situ* hybridization revealed two signals: the expected signal in the 20A region of the X chromosome and additional signal on chromosome 3L, which harbors the endogenous *ubi-p63E* gene (Fig. 3B). The absence of signals in other sites indicates that 20A-X line does not harbor additional transgene insertions at other genomic regions. To further validate these findings, we performed whole-genome sequencing and searched for reads corresponding to junctions between the reporter and genomic sequences. We identified multiple reads corresponding to the two expected flanking regions in the 20A cluster, while no additional insertions were identified, corroborating results of the chromosome hybridization (Fig. S3B). Finally, we employed CRISPR/Cas9 to generate a deletion that removes sequences flanking the insertion site in the 20A-X line. We verified the deletion and concomitant loss of the reporter sequence from the genome by genomic qPCR and loss of GFP expression (Fig. 3C). Thus, the 20A-X line contains a reporter insertion in the single genomic site in the same position as other 20A lines.

**Figure 3.**
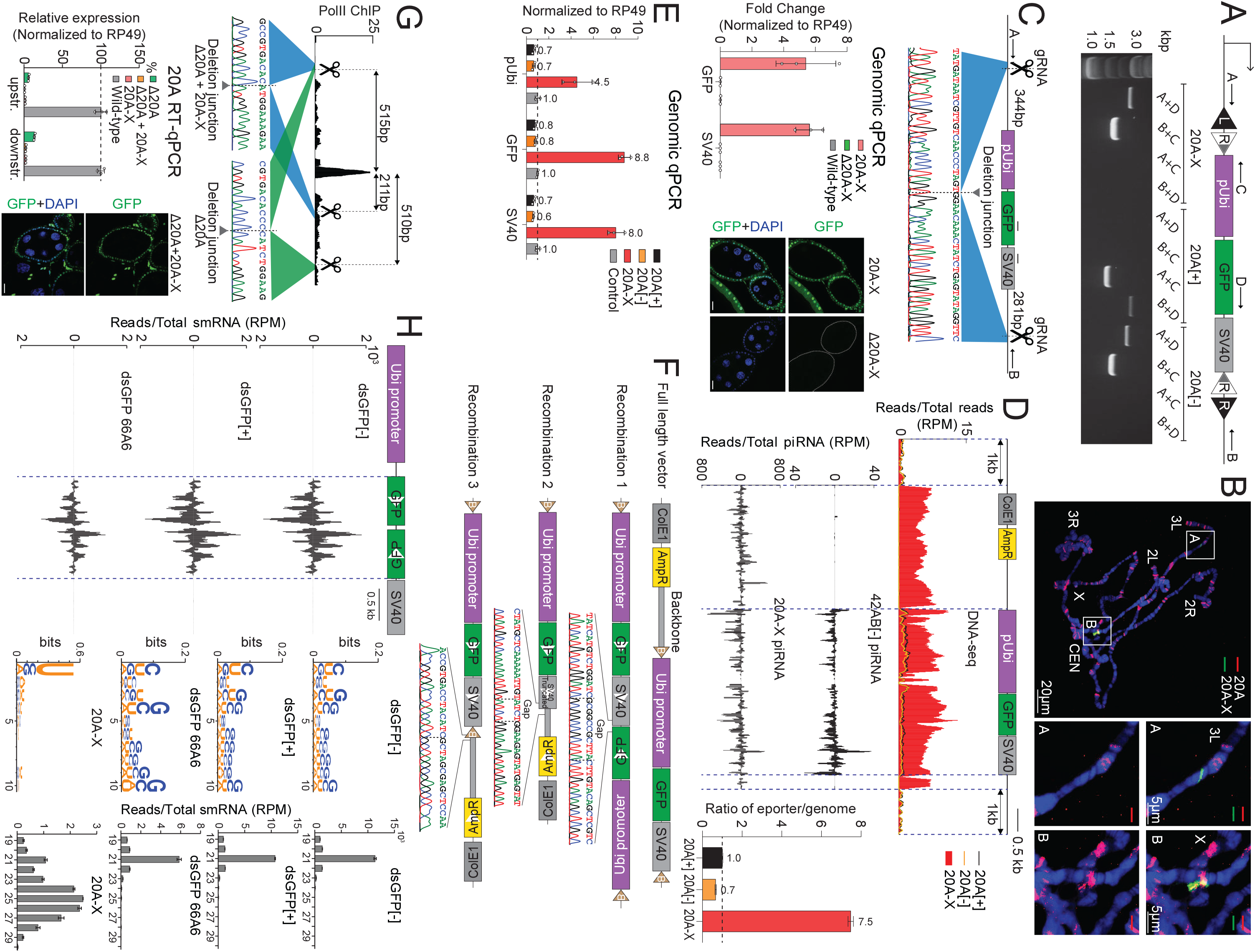
The 20A-X locus contains rearranged, multi-copy reporter sequences. **(A) Determining reporter orientation by genomic PCR**. Positions of the primers are shown. 20A-X has the same minus-strand orientation as the 20A[-] reporter. **(B) 20A-X is located in a single site in the 20A region**. DNA FISH on salivary gland polytene chromosomes was performed using probes against the Ubi-GFP reporter (green, Cy488) and the 20A cluster (red, Cy594). Probes against the reporter detected two locations: one co-localizes with the 20A cluster signal on the X chromosome, the other signal is localized on chromosome 3L where the native ubiquitin gene (*Ubi-p63E*) resides (chr3L:3899259-3903184, dm6). **(C) Verification of the 20A-X insertion site by CRISPR deletion**. The region of the 20A cluster that includes the reporter insertion site was deleted in the 20A-X line using CRISPR/Cas9. Shown are positions of guide RNAs and primers used to verify the deletion using genomic PCR and Sanger sequencing (top). Detection of reporter sequences using qPCR of genomic DNA (left) shows absence of reporter sequences in flies with the deletion. No GFP expression is detected in flies with the deletion (right). Scale bar is 20µm. **(D) 20A-X includes plasmid backbone sequence** (Top) Profiles of whole-genome DNA-seq reads over the sequence of the plasmid used *for* RMCE and the flanking 1 kB genomic sequences in flies with different 20A reporters. 20A-X, but not 20A[+] and 20A[-] flies harbor plasmid backbone sequence, which is normally not integrated during RMCE. Bar graph shows the ratio of reads derived from the reporter to reads from flanking genome sequences in different reporter lines. (Bottom) piRNA profiles over the reporter sequence in 20A-X and 42AB reporter flies. Note that 20A-X generates more piRNA even after normalization to sequence copy number. Profiles represent a mean of two biological replicates and were normalized to total number of piRNA reads in each library. **(E) 20A-X contains multiple copies of reporter sequence**. Different portions of the reporter sequence were measured by genomic qPCR in flies with 20A reporters as well as a control line in which the reporter was integrated into non-cluster region (chr 3L: 7,575,013, dm6). qPCR values were normalized to the rp49 gene region. Fold-differences compared to homozygous flies with the control reporter are indicated above the bars. Error bars indicate standard deviation of three biological replicates. **(F) Reporter sequence rearrangements in 20A-X**. Three abnormal sequence junctions were detected in the 20A-X sequence by DNA-seq, Sanger sequencing PCR-amplified genomic DNA. **(G) Deletion of the 20A cluster promoter does not affect activity of 20A-X**. The putative promoter of the 20A cluster (determined by a prominent Pol II ChIP-seq peak) was deleted using CRISPR/Cas9 in wild-type and 20A-X flies. RT-qPCR shows dramatic reduction of 20A cluster transcripts in flies with the deletion, indicating that it disrupts the cluster promoter. RT-qPCR primers were positioned 2030 and 3769 bp downstream of the putative promoter (upstream and downstream of the reporter insertion, respectively). Error bars indicate standard deviation of three biological replicates. GFP remained repressed in the germline of 20A-X flies upon promoter deletion, indicating that piRNA-mediated silencing remains unaffected. Scale bar is 20µm. **(H) Inverted repeat reporters generate abundant endo-siRNA, but not piRNA**. Scheme of the inverted repeat dsGFP reporter. The dsGFP reporter harbors the same sequence fragments as other reporters, however, the GFP sequence forms an inverted repeat that generates hairpin dsRNA after transcription from the ubiquitin promoter. Shown are small RNA profiles (19-30 nt) along the reporter sequence in ovaries of flies with reporter integration into the 20A cluster in both orientations as well as integration into the control non-cluster region (66A6, chr 3L: 7575013, dm6). Shown on the right are size distributions of reporter mapping small RNAs and nucleotide composition of 23-29 nt (piRNA size range) RNAs. Size profile and nucleotide bias of 20A-X small RNAs are shown for comparison. dsGFP reporters generate abundant endo-siRNAs exclusively from the inverted repeat sequence. The miniscule amount of 23-29nt small RNA generated from dsGFP reporters does not show a 1U-bias expected from genuine piRNAs.

We employed whole-genome DNA-seq and qPCR to analyze reporter copy number in the genome, which showed that while other 20A reporters harbor a single copy of the reporter sequence, 20A-X contains 10 copies (Fig. 3D, E, S3D). In addition, both approaches revealed that, unlike 20A[+] and 20[-] lines, the 20A-X insertion contains plasmid backbone sequence that is used for recombinase-mediated cassette exchange and is normally removed during this process (Fig. 1A, 3E, S3D). Furthermore, we also found multiple DNA-seq reads indicating three unexpected junctions: (1) between the SV40 3’ UTR and the GFP sequence, (2) between the SV40 3’ UTR and the plasmid backbone and (3) between the attB site and the plasmid backbone (Fig. 3F). We confirmed all junctions by genomic PCR and Sanger sequencing. As in situ hybridization on polytene chromosomes, DNA-seq and CRISPR/Cas9 deletion all indicate a single insertion site in the genome, all reporter copies are located in a single genomic site. Taken together, our results indicate that the 20A-X line contain multiple, rearranged copies of the reporter sequence in a single site within the 20A cluster. Together, analysis of 20A-X indicates that, unlike single-copy reporters in 20A, repetitive sequences inserted in the same site generate piRNAs that induce repression in the germline.

To explore if the ‘host’ 20A cluster is required for 20A-X to function as dual-strand piRNA cluster we employed CRISPR/Cas9 to delete the promoter of the 20A cluster (Fig. 3G). In wild-type flies, deletion of the putative promoter eliminated expression of long RNA (piRNA precursors) from the 20A cluster, indicating that deletion disrupts its function. However, deletion of the 20A cluster promoter in 20A-X flies did not release reporter silencing (Fig. 3G), indicating that piRNA-dependent repression of the 20A-X locus does not require activity of the ‘host’ 20A cluster.

20A-X is inserted in the same site as 20A[+] and 20[-] reporters but differs in its repetitive nature as well as its structural rearrangements. Particularly, 20A-X harbors inversions that are expected to generate dsRNA upon their transcription. To explore if the presence of transcribed inverted repeats is sufficient to create a functional dual-strand piRNA cluster, we generated a dsGFP reporter, which consists of the same sequence fragments as the original reporter, but harbors inverted GFP sequences that would form dsRNA upon transcription (Fig. 3H). We obtained flies with insertion of the dsGFP reporter into the same site as other 20A reporters as well as in a control (non-piRNA cluster) region of the genome (chr 3L: 7,575,013). Analysis of small RNAs in ovaries of flies carrying dsGFP constructs revealed the presence of abundant 21 nt siRNAs generated from the inverted GFP repeat, but no other portion of the construct (Fig. 3H). In contrast to 20A-X, dsGFP insertions produce only miniscule amount of larger RNA species and these RNAs lack a U-bias. There were no significant differences between small RNAs generated from the dsGFP reporter inserted into the 20A cluster in either orientation and in the non-cluster control genomic region. Overall, these results shows that inverted repeats trigger generation of siRNAs, but not piRNAs, indicating that the presence of inverted repeats might be necessary, but not sufficient to make 20A-X an active dual-strand piRNA cluster and that the multi-copy nature and/or the extended lengths of the locus might play a critical role.

### piRNA-induced repression of the 20A-X reporter depends on maternal transmission of cognate piRNAs

Previously we and others have shown that the activity of artificial dual-strand piRNA clusters in the progeny requires cytoplasmic inheritance of piRNAs from the mother(de Vanssay et al., 2012; Hermant et al., 2015; Le Thomas et al., 2014b), and proposed that all dual-strand clusters might depend on trans-generationally inherited piRNAs to maintain their activity(de Vanssay et al., 2012; Le Thomas et al., 2014b). Therefore, we explored expression of the 20A-X reporter in the progeny after paternal and maternal inheritance. Flies that inherited the 20A-X insertion from their mothers (maternal transmission) showed – similar to their mothers – GFP expression in follicular cells, but strong GFP repression in the germline with sense reporter RNA restricted to the nucleus. In contrast, females that inherited the 20A-X reporter from their fathers (paternal transmission) had robust GFP protein expression and cytoplasmic localization of GFP mRNA in the germline (Fig. 4A). We also observed GFP repression in the germline of males that inherited the 20A-X locus maternally, but not when they inherited it paternally. In agreement with FISH and IF results, RT-qPCR showed ∼5-fold increase of GFP RNA after paternal transmission (Fig. 4B). Thus, repression of the 20A-X reporter in both in the male and female germline requires maternal inheritance of the reporter.

**Figure 4.**
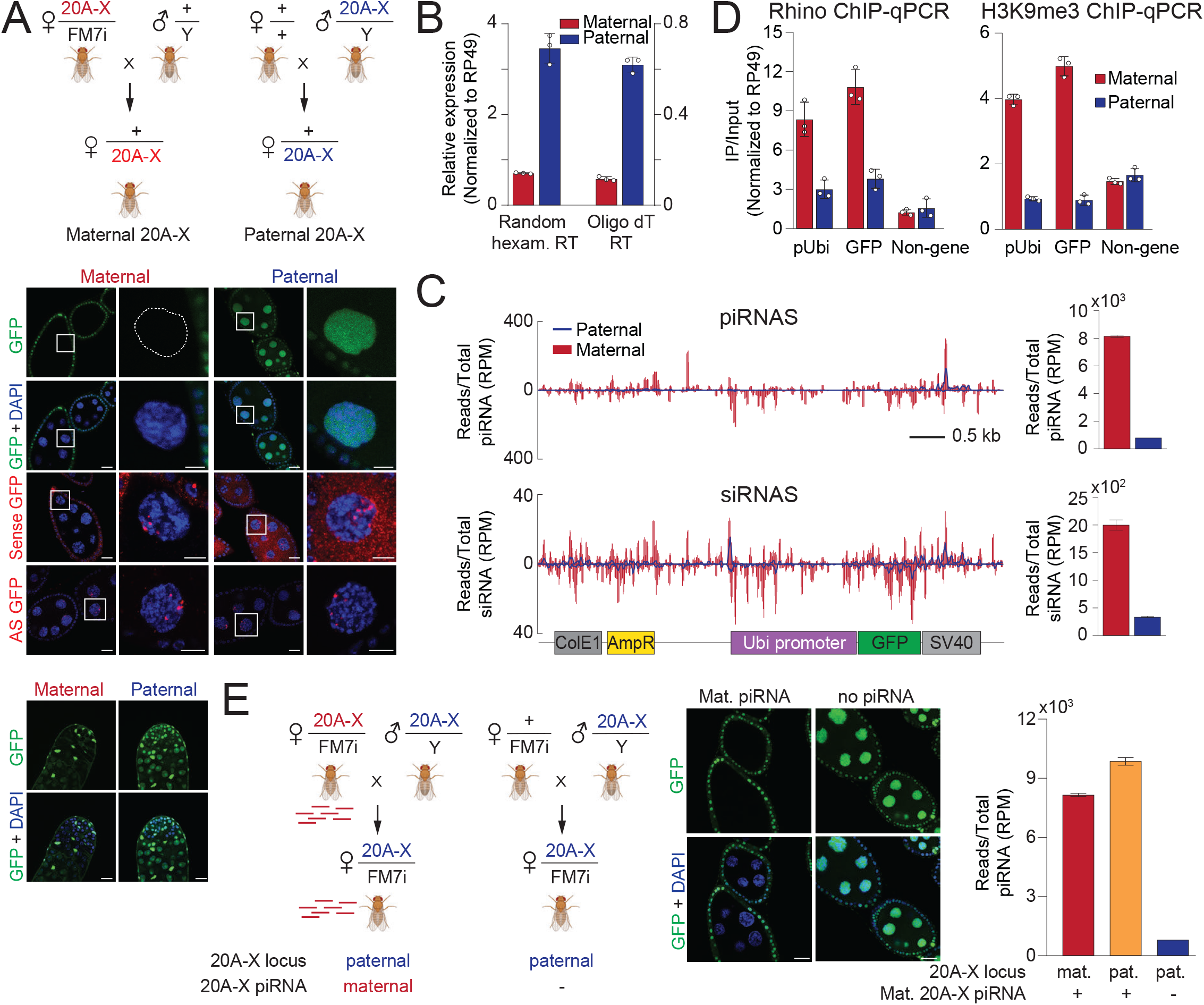
piRNA-induced repression of the 20A-X reporter depends on maternal transmission of cognate piRNAs. **(A) 20A-X repression is released after paternal transmission**. Scheme of crosses to test effects of maternal and paternal transmission of 20A-X. Note that the genotype of the progeny from the two crosses are identical. Shown are GFP protein and sense and antisense GFP RNA expression in ovaries and GFP protein expression in testes of the progeny. Scale bar is 20µm and 2µm for egg chamber and single nurse cell nuclei, respectively, and 20µm for testis. **(B) GFP RNA expression increases after paternal transmission of 20A-X**. RT-qPCR of GFP RNA in ovaries of progenies from the two crosses in (A) were performed with random hexamer and oligo dT primers and normalized to rp49 mRNA level. Error bars indicate the standard deviation of three biological replicates. **(C) 20A-X piRNA level drops after paternal inheritance**. Shown are piRNA and siRNA profiles along the reporter sequence in ovaries of progeny from the two crosses shown in (A). Bar graphs on the right show number of piRNA and siRNA reads mapping to the reporter normalized to total piRNA and siRNA reads, respectively. Error bars indicate standard deviation of two biological replicates. **(D) Rhino and H3K9me3 are lost on 20A-X chromatin after paternal transmission**. The levels of Rhino and H3K9me3 on chromatin were measured by ChIP-qPCR using primers against the pUbi and GFP regions as well as a control non-cluster region (chr 2L: 968,088 – 968,187, dm6) and normalized to the *rp49* locus. H3K9me3 and Rhino levels drop after paternal transmission to levels similar to that of the control region. Error bars indicate the standard deviation of three biological replicates. **(E) Cytoplasmic piRNA inheritance is sufficient for repression of paternally transmitted 20A-X**. Scheme of crosses to test the effect of maternal cytoplasmic piRNA inheritance on repression of paternally transmitted 20A-X. Progeny of both crosses inherit the 20A-X locus from their fathers and are genetically identical. The only difference is cytoplasmic inheritance of small 20A-X-derived RNAs from their mothers, which is sufficient to restore both GFP repression and piRNA generation in the germline of the progeny. Number of piRNA reads mapping to the reporter was normalized to total piRNA reads mapping to the genome in each library. Error bars indicate standard deviation of two biological replicates. Scale bar is 20µm.

To understand if derepression of GFP after paternal transmission is caused by changes in piRNA expression, we cloned small RNA libraries from ovaries of progeny that inherited the locus maternally and paternally. Upon paternal transmission, piRNA level was 10.1-fold reduced (Fig. 4C). Thus, piRNA generation from the 20A-X locus requires its maternal inheritance and derepression of GFP upon paternal reporter transmission is explained by the dramatic decrease in reporter-targeting piRNA. Next, we determined enrichment of the H3K9me3 mark and Rhino protein on chromatin of 20A-X reporter in the progeny upon paternal or maternal inheritance of the locus. Both Rhi and H3K9me3 were reduced on 20A-X chromatin after paternal transmission to levels comparable to those detected at the control euchromatic region (Fig. 4D). Thus, loss of piRNA upon paternal transmission correlates with loss of H3K9me3 and Rhino from the 20A-X locus.

Progeny that inherits the 20A-X locus from their mothers receive two distinct contributions. First, the locus itself might have different chromatin imprints when inherited maternally or paternally. Second, mothers deposit piRNAs into the oocyte, while paternal progeny do not inherit piRNA from their fathers(de Vanssay et al., 2012; Hermant et al., 2015; Le Thomas et al., 2014b). Therefore, it is important to discriminate if the parent-of-origin effects of 20A-X depend on inheritance of the genomic locus or cytoplasmic transmission of piRNAs to the next generation. To discriminate between these possibilities, we designed two different crosses: in both crosses the progeny inherited 20A-X paternally, however, in one cross the mothers also carried a copy of the 20A-X locus which, however, was not transmitted to the progeny (Fig. 4E). The presence of the 20A-X locus in mothers caused GFP repression and piRNA generation in the progeny even though the locus itself was not transmitted to the offspring. Indeed, progeny that inherited the 20A-X locus from their fathers but received cognate piRNAs from their mothers had similar level of 20A-X piRNAs as progeny that simply inherited the 20A-X locus maternally. It is worth noting that maternally inherited cognate piRNAs were not able to convert the 20A[+] and 20A[-] loci to piRNA-producing clusters, nor did they change the expression of GFP from these reporters (data not shown). These results indicate that the activity of 20A-X as a piRNA-generating locus requires both the extended, multi-copy nature of the locus and cytoplasmic inheritance of cognate piRNAs through the maternal germline.

### piRNA cluster is established over several generations

The finding that maternal inheritance of piRNAs is required for the activity of the 20A-X locus in the progeny prompted us to re-examine the observation that initially, upon establishment of the 20A-X transgenic flies by RMCE, GFP was expressed in the germline but got repressed in later generations (Fig. 5A). First, we established that the age of flies did not influence GFP silencing, as repression was similar in young (5-days) and old (30-days) females of the 14 months old stock (Fig. 5A). Next, we analyzed small RNA profiles in ovaries of 20A-X flies 3, 11 and 21 months after establishment of the stock. piRNAs and siRNAs derived from 20A-X were already present at 3 months, but their abundance increased 4-fold and 3.7-fold, respectively, by 11 months (Fig. 5B). At 11 months 20A-X piRNAs also showed stronger sign of ping-pong processing as measured by complementary piRNA pairs that overlap by 10nt (Z-scores at 3 months and 11 months were 1.0 and 4.2, respectively) (Fig. 5C, S5A). No further increase in abundance of 20A-X piRNAs and ping-pong processing was observed when comparing 11 and 21 months-old stocks. Thus, transgene-derived piRNA abundance is increasing over multiple generations after transgenesis and this increase correlates with repression of GFP in the germline. The findings that 20A-X requires maternally-supplied piRNAs in order to generate piRNAs suggests that 20A-X was not active as a piRNA cluster in the first generation after establishment of this stock.

**Figure 5.**
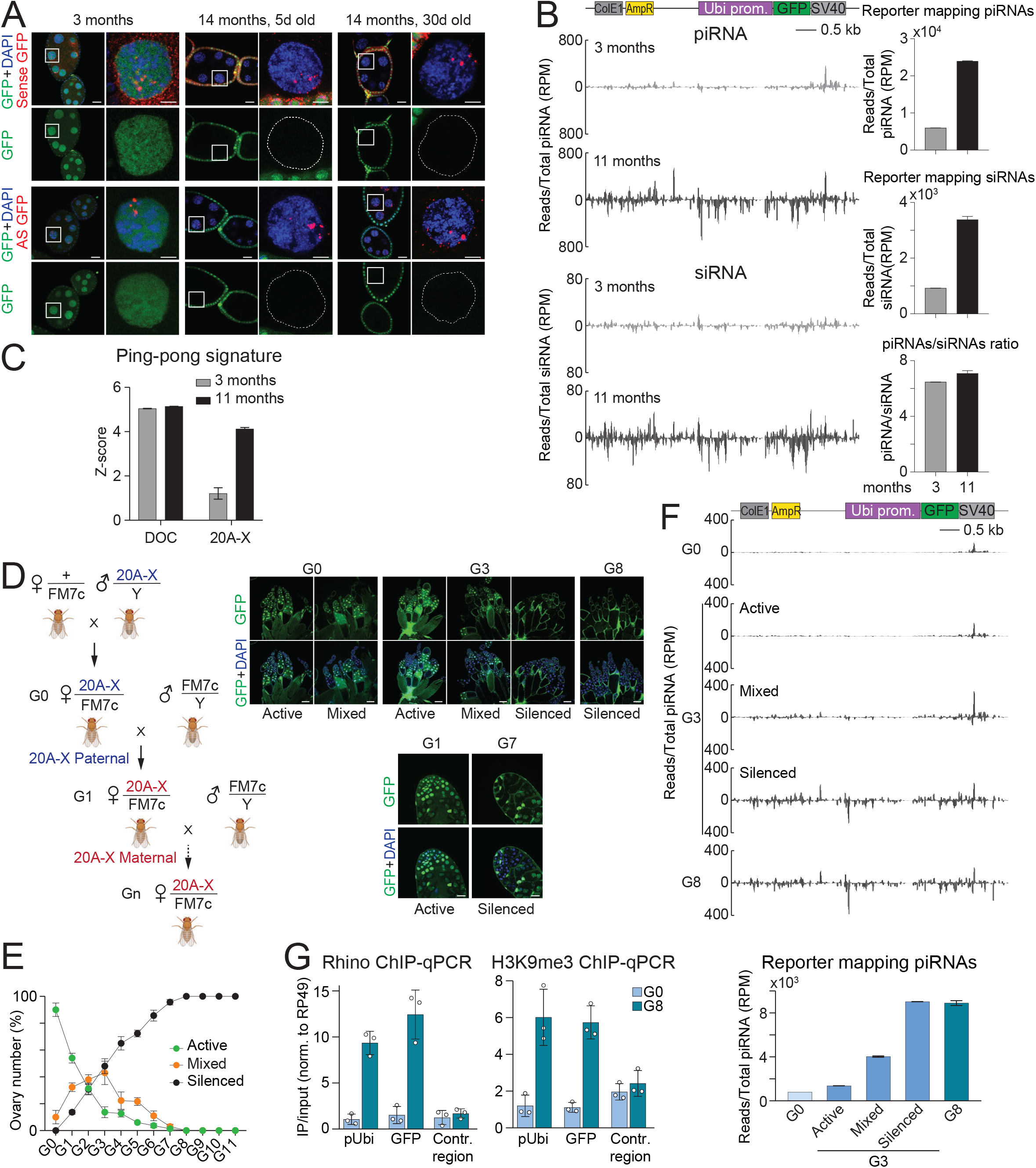
piRNA cluster is established over several generations. **(A) Establishment of 20A-X repression over several generations**. Shown are GFP protein and RNA expression in ovaries of 20A-X flies 3 and 14 months after establishment of this line by RMCE. Young (5 days after hatching) and old (30 days after hatching) flies show no difference in expression. Scale bar is 20µm and 2µm for egg chamber and single nurse cell nuclei, respectively. **(B) 20A-X piRNA level increases over several generations**. Shown are ovarian reporter-mapping piRNA and siRNA profiles 3 and 11 months after establishing the 20A-X line. Bar graph shows the levels of 20A-X piRNAs and siRNAs normalized to total number of piRNA and siRNA reads in each library. Error bars indicate standard deviation of two biological replicates. **(C) Ping-pong signature of 20A-X-mapping piRNAs increases over several generations**. Shown are Z-scores indicating ping-pong signature (10 nt distance between 5’ ends of complementary piRNAs) of 20A-X piRNAs. Z-scores of DOC transposon piRNA are shown for comparison. Error bars indicate standard deviation of two biological replicates. **(D) Recovery of 20A-X repression after paternal transmission**. Left: Scheme of crosses to monitor 20A-X after its paternal transmission. After paternal transmission in the first cross (G0), 20A-X is inherited maternally in each subsequent generation (G1-G8). Right: Expression of GFP in selected generations in ovaries and testes. Several generations (G1-G5) show variable GFP expression between individual flies and within each fly ovary. Scale bar is 100µm and 20µm for ovary and testis, respectively. **(E) Accumulation of 20A-X repression over several generations**. In each fly germline GFP expression was determined and assigned one of three values: ‘silenced’ indicates complete lack of expression, ‘active’ indicate expression in the majority of germline nuclei, while ‘mixed’ indicate variable expression between individual egg chambers within the same ovary. Plotted is the fraction of ovaries with corresponding expression pattern in each generation from G0 to G11. The experiment was repeated three times and 100 ovaries were counted in each generation in each replica. Error bars indicate the standard deviation of three biological replicates. **(F) Accumulation of 20A-X piRNAs over several generations**. Shown are profiles of 20A-X piRNAs in different generations. In G3, ovaries were separated in three groups according to GFP expression as described in (E) and for each group independent small RNA libraries were prepared. Bar graph (bottom) shows 20A-X piRNA levels normalized to total piRNA reads in each library. Error bars indicate standard deviation of two biological replicates. **(G) Accumulation of Rhino and H3K9me3 on 20A-X chromatin**. Rhino and H3K9me3 levels on chromatin were measured by ChIP-qPCR in ovaries of G0 and G8 generation using primers against the pUbi and GFP regions as well as a control, non-cluster region (chr 2L: 968,088 – 968,187, dm6) and normalized to the *rp49* locus. Error bars indicate the standard deviation of three biological replicates.

The loss of 20A-X’s ability to generate piRNAs upon paternal inheritance provides a unique opportunity to explore *de novo* establishment of a piRNA cluster. After paternal transmission the progeny (G0) generated very few piRNAs and siRNAs with levels similar to those of 20A[+] and 20A[-] flies (after normalization to transgene copy numbers) (Fig. S5C). We monitored whether 20A-X can recover its ability to generate piRNAs in future generations upon continuous maternal transmission (Fig. 5D). While no GFP repression occurred in the germline of G0, each subsequent generation showed decreased GFP expression, until complete repression was observed in G8 (Fig. 5D, S5B). Establishment of GFP repression over multiple generations also occurred in the male germline. Interestingly, ovaries of flies of intermediate generations (G2-G5) showed large variation in the extent of repression between individual egg chambers (Fig. 5D, E, S5B). For example, in G2 almost equal fractions of flies showed normal expression, complete silencing or a mixed phenotype (Fig. 5E).

We profiled small RNAs in ovaries of G3 flies after separating them into three groups (active, mixed and silenced) based on GFP expression as well as from ovaries of G0 and G8 flies. Reporter piRNA levels increased over generations. Importantly, the three groups of G3 ovaries with different levels of GFP repression had proportionally different levels of reporter piRNAs, indicating that repression correlates with piRNA abundance (Fig. 5F).

Finally, we determined enrichment of H3K9me3 mark and Rhino protein on chromatin of the 20A-X reporter in ovaries of G0 and G8 progeny. While Rhi and H3K9me3 were lost after paternal transmission in G0, both Rhi and H3K9me3 were enriched on 20A-X chromatin in G8 (Fig. 5G). Overall, our results indicate that 20A-X gradually establishes its ability to generate piRNAs over eight generations if maternal 20A-X piRNAs are transmitted to the progeny in each generation.

### Maternal siRNA triggers activation of piRNA cluster in the progeny

How loci like 20A-X start to function as dual-strand cluster and generate piRNAs remains unclear. In addition to piRNAs, all studied dual-strand clusters generate siRNA. Although we found that transgenes containing simple inverted repeats produce exclusively siRNAs and no piRNAs, it is still possible that the presence of cognate siRNAs might provide the initial trigger to activate piRNA production.

To test the role of siRNAs in piRNA cluster activation, we crossed males carrying the 20A-X locus with heterozygous females carrying dsGFP constructs harboring simple inverted repeats that generate siRNAs (Fig. 6A). As seen before, paternal transmission of 20A-X led to release of GFP repression in the germline of the progeny (Fig. 6B). However, GFP remained repressed in progeny that carried maternally-inherited dsGFP constructs. Remarkably, a similar level of GFP repression was also observed in sibling progeny that did not inherit the dsGFP construct from their mothers, indicating that the presence of cognate siRNAs in the mothers was sufficient to activate repression in progeny (Fig. 6B).

**Figure 6.**
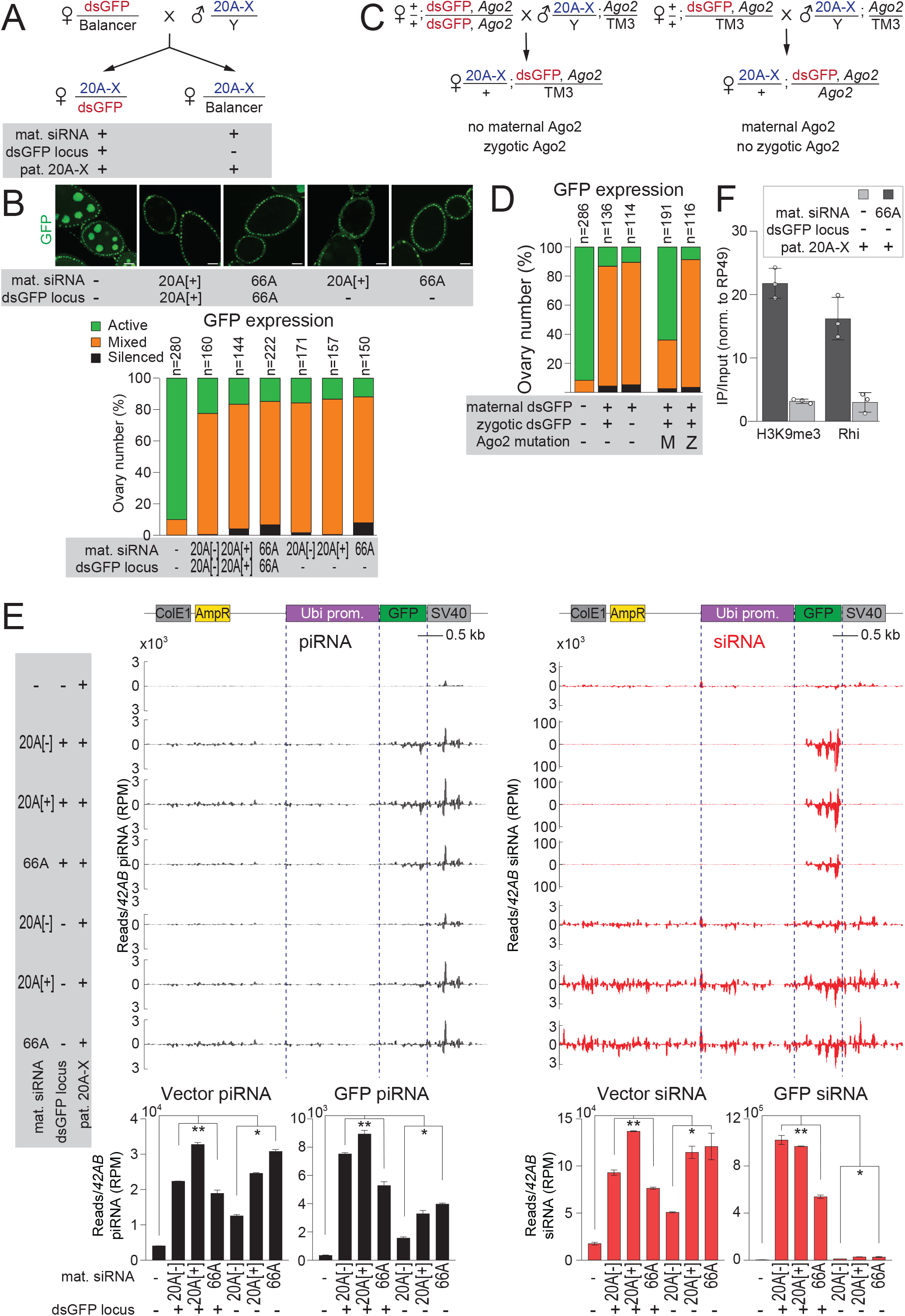
Cytoplasmic inheritance of siRNAs activates piRNA biogenesis in the progeny. **(A) Crossing scheme to test the role of siRNAs in triggering 20A-X repression**. The 20A-X reporter is inherited from the father, while mothers harbor an inverted repeat dsGFP construct that generates siRNA. Progeny that inherited the dsGFP locus and those that did not were compared. **(B) Cytoplasmic inheritance of siRNAs triggers 20A-X repression in the progeny**. GFP expression in progenies of each genotype was assessed by fluorescent microscopy (top) and assigned one of the three values as described in Fig. 5E (bottom). Two independent dsGFP loci in 20A and in a locus in band 66A were analyzed. Maternal siRNA triggered repression of 20A-X in the progeny independently of inheritance of siRNA-generating locus or the genomic position of the siRNA-generating locus. Scale bar is 20µm. N indicates the number of ovaries analyzed for each genotype. GFP expression in progenies of each genotype was assessed by fluorescent microscopy. **(C) Crossing scheme to test the role of Ago2 in 20A-X repression**. Crosses are similar to crosses shown in **(A)** except of the presence Ago2 mutation either in mothers or the progeny. **(D) Triggering of 20A-X repression by trans-generational siRNAs depends on a functional siRNA pathway in the mothers, but not in the progeny**. Analysis of GFP expression in progenies of crosses shown in (C). **(E) Cytoplasmic inheritance of siRNAs activates piRNA production in the progeny**. Shown are profiles of piRNAs and siRNAs mapping to the reporter in ovaries of progenies of crosses shown in (A). Bar graph shows 20A-X piRNA and siRNA levels normalized to those generated from the *42AB* cluster. Error bars indicate standard deviation of two biological replicates. Statistical significance is estimated by two-tailed Student’s t-test; *p<0.05, **p<0.01. **(F) Cytoplasmic siRNA inheritance is required for accumulation of H3K9me3 and for Rhino recruitment**. Rhino and H3K9me3 level on chromatin of 20A-X was measured by ChIP-qPCR in progenies of crosses shown in (A). ChIP signal in 20A-X is normalized to the *rp49* locus. Error bars indicate the standard deviation of three biological replicates.

To further explore the role of siRNAs, we abrogated the siRNA pathway either in the mothers or in the progeny using Ago2 mutation (Fig. 6C). In flies, Ago2 is required for the stability and function of siRNAs and its mutation completely disrupts the siRNA pathway(Okamura et al., 2004). GFP repression was strongly disrupted in the progeny of Ago2-deficient mothers, confirming that it requires trans-generational cytoplasmic transmission of siRNA (Fig. 6D). In contrast, Ago2-deficient progeny that inherited siRNA from their heterozygous mothers show strong GFP repression. Taken together, these results indicate that initiation of 20A-X repression in the progeny requires trans-generational inheritance of cytoplasmic siRNAs, while the siRNA pathway is dispensable for maintenance of the repression.

To further analyze the effect of siRNA on activation of 20A-X we cloned and sequenced small RNAs. As expected, progeny that inherited the dsGFP construct had high level of siRNAs targeting GFP (>100-fold increase compared to progeny with only paternal 20A-X) (Fig. 6E). These abundant siRNAs were restricted to the GFP sequence, which forms inverted repeats in the dsGFP construct. Interestingly, sibling progeny that inherited the balancer chromosome instead of dsGFP also showed moderate (3∼7-fold) increase in GFP siRNA level compared to flies with paternal 20A-X only. Remarkably, both the progeny that inherited the dsGFP construct and their siblings that inherited the balancer chromosome had elevated levels of piRNA mapping to the 20A-X reporter (Fig. 6E). piRNAs mapping to the GFP sequence were 15-26-fold more abundant in progeny that inherited the dsGFP constructs and 5-12-fold more in progeny with the balancer chromosome when compared to flies that only had the paternal 20A-X. However, even more remarkably, both progenies that inherited dsGFP and those with the balancer chromosome had similar, 3 to 8-fold elevated levels of piRNA produced from regions of 20A-X that are not targeted by GFP siRNAs. This means that maternally contributed GFP siRNAs that target a portion of the 20A-X locus were sufficient to induce piRNA generation from the entire 20A-X locus in the progeny. Furthermore, using ChIP-qPCR we found that cytoplasmic inheritance of GFP siRNAs from the mother was sufficient to trigger accumulation of H3K9me3 and Rhi on chromatin of paternally-inherited 20A-X (Fig. 6F). These results suggest that siRNAs are able to provide the initial trigger that converts the 20A-X locus into a dual-strand piRNA cluster.

## Discussion

### siRNAs can provide initial trigger to activate piRNA biogenesis

As a system that protects the genome against selfish genetic elements, the piRNA pathway has to be able to adapt to target new invader elements. Previous studies have revealed mechanisms to store information about genome invaders in piRNA clusters and to maintain piRNA biogenesis through a feed-forward loop that involves trans-generational cytoplasmic transmission of piRNAs. These inherited piRNAs guide deposition of the RDC chromatin complex on clusters, which is required for efficient piRNA expression in the progeny. However, the question of how the pathway adapts to new TEs and starts creating piRNAs against novel threats remained unresolved. One possibility for adaptation is integration of new transposons into pre-existing piRNA clusters, which would lead to generation of novel piRNAs, a process that has been modeled experimentally (Le Thomas et al., 2014b; Muerdter et al., 2012) and observed naturally (Khurana et al., 2011; Zhang et al., 2020). However, other studies suggest that entire new piRNA-generating regions can arise in evolution, providing another mechanism for acquiring immunity against novel elements. Indeed, some piRNA clusters are active only in one but not other *D. virilis* strains (Le Thomas et al., 2014a), suggesting their recent formation. Recent comprehensive evolutionary analysis showed that piRNA cluster regions are extremely labile in *Drosophila* evolution, suggesting frequent acquisition and loss of piRNA clusters (Gebert et al., 2021). Finally, spontaneous formation of novel piRNA clusters from transgenic sequences has been observed (de Vanssay et al., 2012). Though these findings suggested that new piRNA-producing genomic regions that contain no sequence homology to pre-existing piRNAs can arise, the conceptual framework to explain this process was lacking.

The finding that siRNAs can activate piRNA biogenesis provides an explanation for how immunity to new transposons can be established through initial detection of new element by the siRNA pathway, which then triggers a stable piRNA response. In contrast to the piRNA pathway that relies on genetic and epigenetic memory – in form of active piRNA clusters – to recognize its targets, the siRNA pathway uses a simple rule for self/non-self discrimination. Unlike normal genes, transposons often generate both sense and antisense transcripts that form dsRNAs, which are recognized by Dicer and processed into siRNAs. Indeed, in addition to sense transcription from their own promoters, transposons are often transcribed in antisense orientation from host genes’ promoters, for example if a TE is inserted into an intron or the 3’UTR in opposite orientation than the host gene orientation. In addition, recursive insertion of transposons into each other creates inverted repeats that generate hairpin RNAs. The presence of sense and antisense transcripts leads to dsRNA and siRNA formation, providing a simple yet efficient mechanism to discriminate mobile genetic elements from host genes. Indeed, the siRNA pathway has well-established functions in recognizing and suppressing both endogenous (transposons) and exogenous (viruses) invader genetic elements in all eukaryotic lineages, especially in TE-rich plant genomes. In contrast, the piRNA pathway is restricted to Metazoa, suggesting that it is a more recent evolutionary innovation. As both pathways target foreign genetic elements, siRNAs provide an ideal signal to activate piRNA biogenesis against novel invaders. Activation of piRNA response by siRNAs can be compared to stimulation of the robust and long-lasting adaptive immune response by the first-line innate immune systems (Medzhitov, 2007).

Our results indicate that siRNAs are important to jump-start piRNA cluster activity, but are dispensable later on. Indeed, cytoplasmically-inherited siRNAs are sufficient to trigger piRNA biogenesis in the progeny, while the zygotic siRNA pathway is completely dispensable for this process (Fig. 6). Consistent with siRNAs being dispensable for maintenance of piRNA biogenesis, a previous study showed that the siRNAs are not required for the activity of an artificial piRNA cluster (T1/BX2) (de Vanssay et al., 2012). On the other hand, increase in the level of antisense transcripts was proposed to be linked to spontaneous activation of BX2 (Casier et al., 2019), suggesting a general role of antisense RNA (and hence the siRNA pathway) in activation of piRNA clusters. Overall, our results and the previous findings suggest a two-step model of cluster activation (Fig. 7): during the first step siRNAs activate piRNA generation from cognate genomic regions, while during the second step continuous generation and maternal inheritance of piRNAs reinforces piRNA biogenesis making siRNAs dispensable for cluster maintenance.

**Figure 7.**
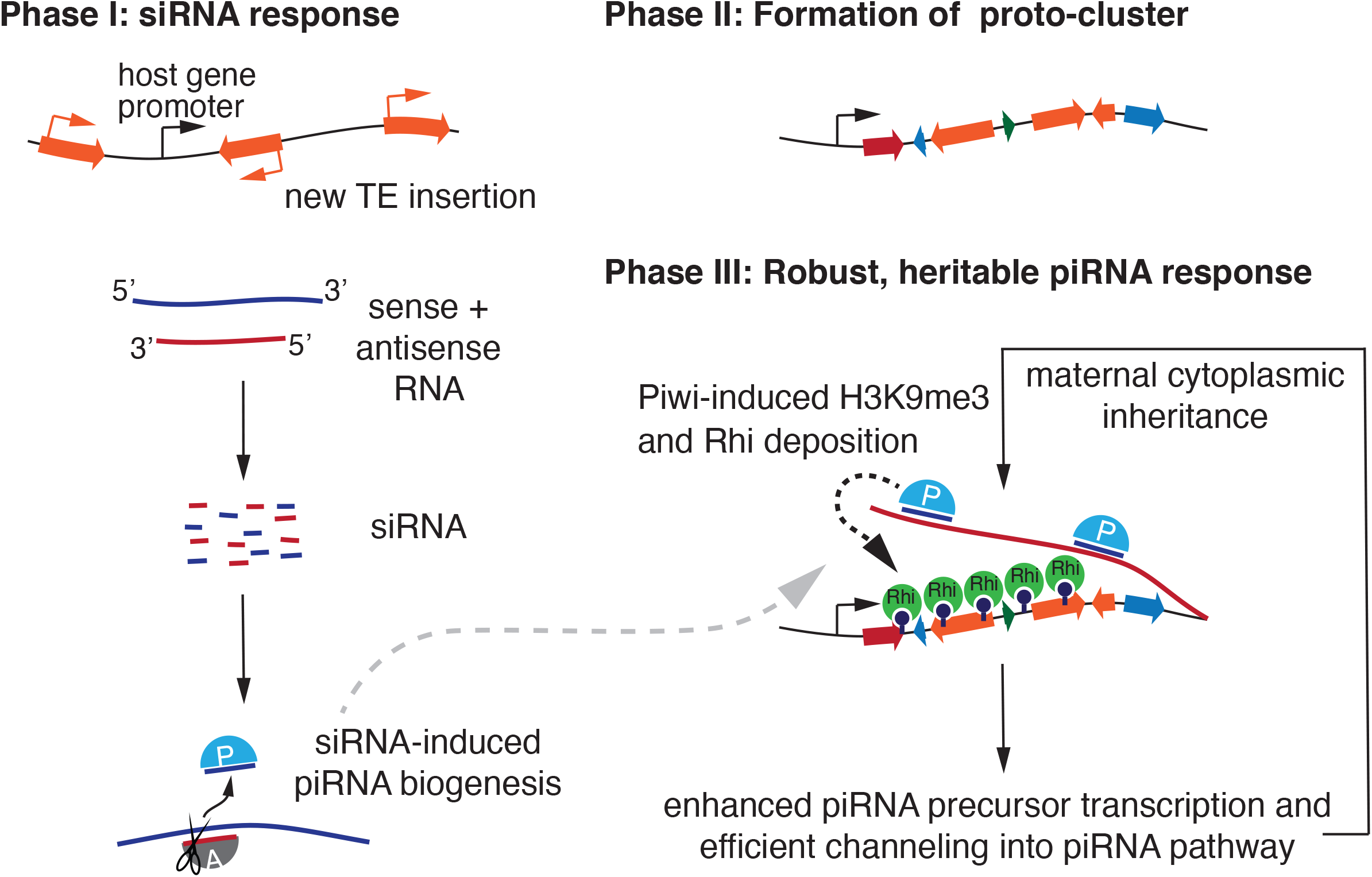
The model for siRNA-triggered activation of piRNA immunity.

The precise molecular mechanism by which siRNAs trigger piRNA biogenesis remains to be understood. In yeast, siRNAs guide a methyltransferase complex to their genomic targets leading to H3K9me3 deposition on chromatin (Verdel et al., 2004). In *Drosophila*, this modification is required for deposition of the RDC complex and thus robust piRNA biogenesis (Le Thomas et al., 2014b; Mohn et al., 2014; Yu et al., 2015a; Zhang et al., 2014). However, in flies siRNA repression seems to be restricted to target RNA cleavage in the cytoplasm, suggesting that they induce piRNA biogenesis through a different mechanism. The cleavage of complementary transcripts by siRNAs creates aberrant RNAs with 5’-monophosohorylated ends, which are good substrates for the cytoplasmic piRNA processing machinery. Thus, we propose that siRNA-induced cleavage of complementary transcripts generates substrates for piRNA processing. Cytoplasmic piRNA processing, in turn, generates piRNAs that are loaded into the three piwi proteins, including the nuclear Piwi protein (Huang et al., 2017; Pandey et al., 2017; Rogers et al., 2017). As the nuclear Piwi/piRNA complex guides establishment of the H3K9me3 mark, this model explains how cytoplasmic siRNAs are capable of inducing chromatin changes that are associated with piRNA cluster activation.

### Genomic requirements for piRNA cluster function

While our results indicate that siRNAs can trigger piRNA biogenesis from cognate genomic regions, we and others also found that simple inverted repeats generate exclusively siRNAs and not piRNAs. Thus, not every region that generates self-targeting siRNAs turns into a piRNA cluster, indicating that siRNAs might be necessary but not sufficient to convert a region into a piRNA cluster. Although 20A-X is composed of the same sequences as the simple inverted repeat construct, it contains ∼ 10 copies of the transgenic sequence. The only other artificial piRNA clusters described, T1/BX2, also contain multiple tandem sequences. Thus, the extended length and repetitive nature seems to be important for *de novo* establishment of clusters. These features might be linked to the important role that chromatin organization plays in piRNA cluster function. Specifically, the extended repetitive organization might be required for maintenance of the RDC chromatin compartment, which is essential for transcription and post-transcriptional processing of piRNA precursors(Andersen et al., 2017; Chen et al., 2016; ElMaghraby et al., 2019; Hur et al., 2016; Klattenhoff et al., 2009; Mohn et al., 2014; Zhang et al., 2014). piRNA clusters are enriched in the heterochromatic H3K9me3 mark and the RDC complex that binds this mark(Klattenhoff et al., 2009; Le Thomas et al., 2014b; Mohn et al., 2014; Yu et al., 2015a; Zhang et al., 2014). Rhi is a paralog of HP1 that forms ‘classic’ heterochromatin domains, which depend on cooperative interactions between multiple HP1 molecules and associated proteins (through interactions between HP1 dimers bound to neighboring nucleosomes)(Le Thomas et al., 2014b; Yu et al., 2015a; Yu et al., 2018). Accordingly, extended length might be required for formation of a stable HP1 chromatin compartment. Tandem repeats seem to be particularly prone to formation of heterochromatin, though the underlying mechanisms is not completely clear(Dorer and Henikoff, 1994). Similar to HP1, Rhi is capable of self-interactions through its chromo and chromo-shadow domains and these interactions are required for formation of RDC compartments and the function of piRNA clusters(Le Thomas et al., 2014b; Yu et al., 2015a). Thus, extended length and tandem repeats might be necessary to form a stable RDC chromatin compartment in the nucleus. Establishment of such a region that is capable of maintaining RDC-rich heterochromatin might be the first step in developing piRNA immunity to a new element (Fig. 7).

## Acknowledgements

We thank members of the Aravin and Fejes Toth labs for discussion and comments. We thank Julius Brennecke for providing the Rhino antibody. We thank Igor Antoshechkin (Caltech) for help with sequencing. This work was supported by grants from the National Institutes of Health (R01 GM097363) and by the HHMI Faculty Scholar Award to AAA.

## Supplementary Material

**Figure S1.**
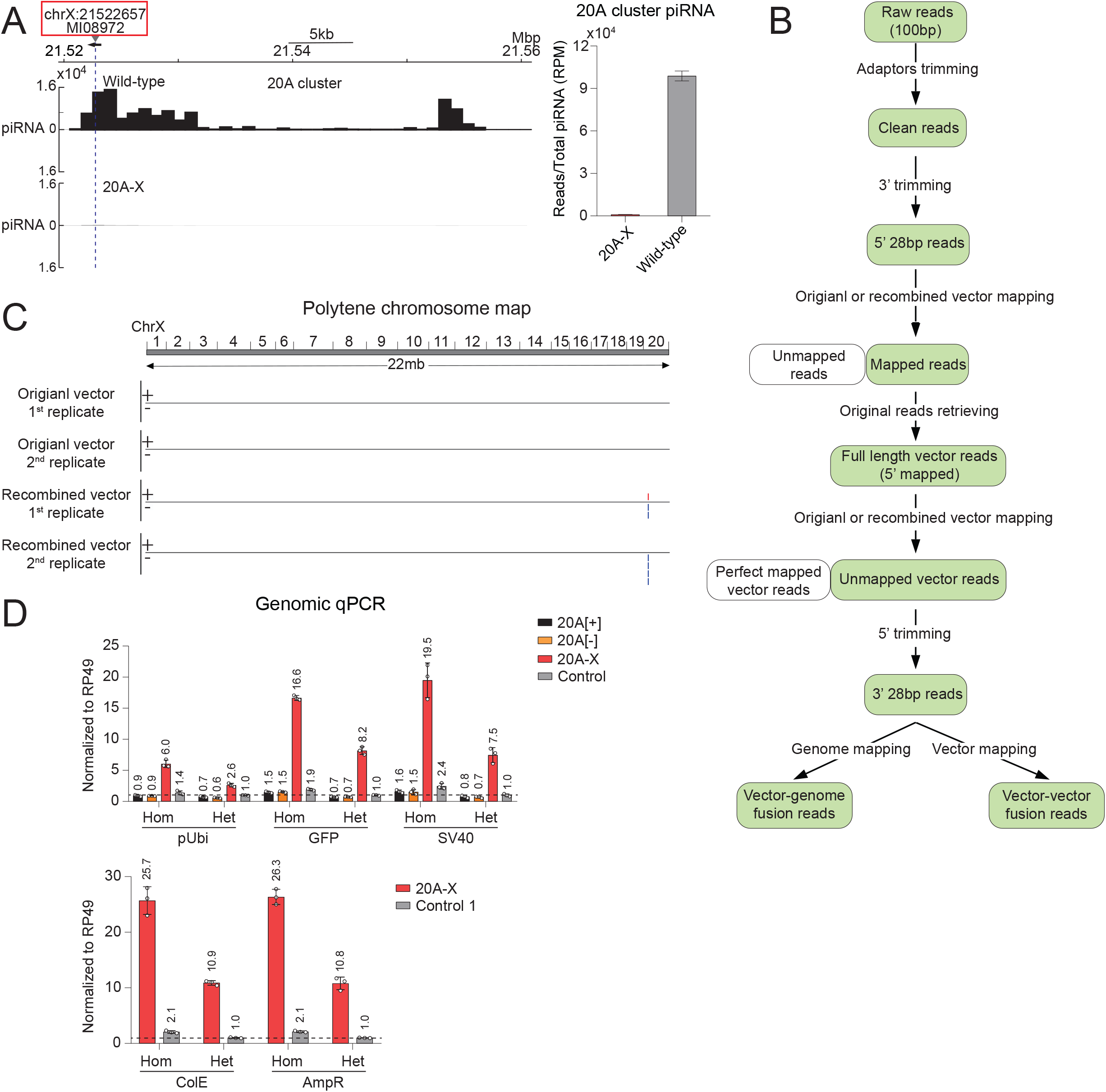
related to Figure 3. Analysis of 20A-X locus organization. **(A)** (Left) Profiles of uniquely-mapping piRNAs of 20A cluster piRNA showed reduced in ovaries of 20A-X flies. Number of piRNA reads mapped to the 20A cluster is normalized to total piRNA read count. (Right) Total piRNA reads uniquely-mapping to 20A cluster were normalized to total piRNA read count. Error bars indicate standard deviation of two biological replicates. **(B)** Pipeline to analyze genomic DNA-seq data to find reporter fusion reads. **(C)** Sequencing of the 20A-X genome revealed a single site of reporter sequence integration in the expected position on the X chromosome. Original vector is the vector prior to recombinase-mediated cassette exchange (including the vector backbone); recombined vector is after recombinase-mediated cassette exchange (without the vector backbone). Analysis was done in two replicates. **(D)** Top: Different portions of the reporter sequence were measured by genomic qPCR in flies with homozygous (without X chromosome balancer) and heterozygous (with X chromosome balancer) 20A reporters as well as control reporter integrated into a non-cluster region (chr3L:7575013, dm6). Bottom: control1 reporter contains a single copy of ColE and AmpR (chr3L:11070538, dm6). The copy number is about two fold more in 20A reporter flies that are homozygous compared to heterozygous flies. qPCR values were normalized to rp49 gene region. Fold-differences compared to heterozygous control reporter are indicated above the bars. Error bars indicate standard deviation of three biological replicates.

**Figure S2.**
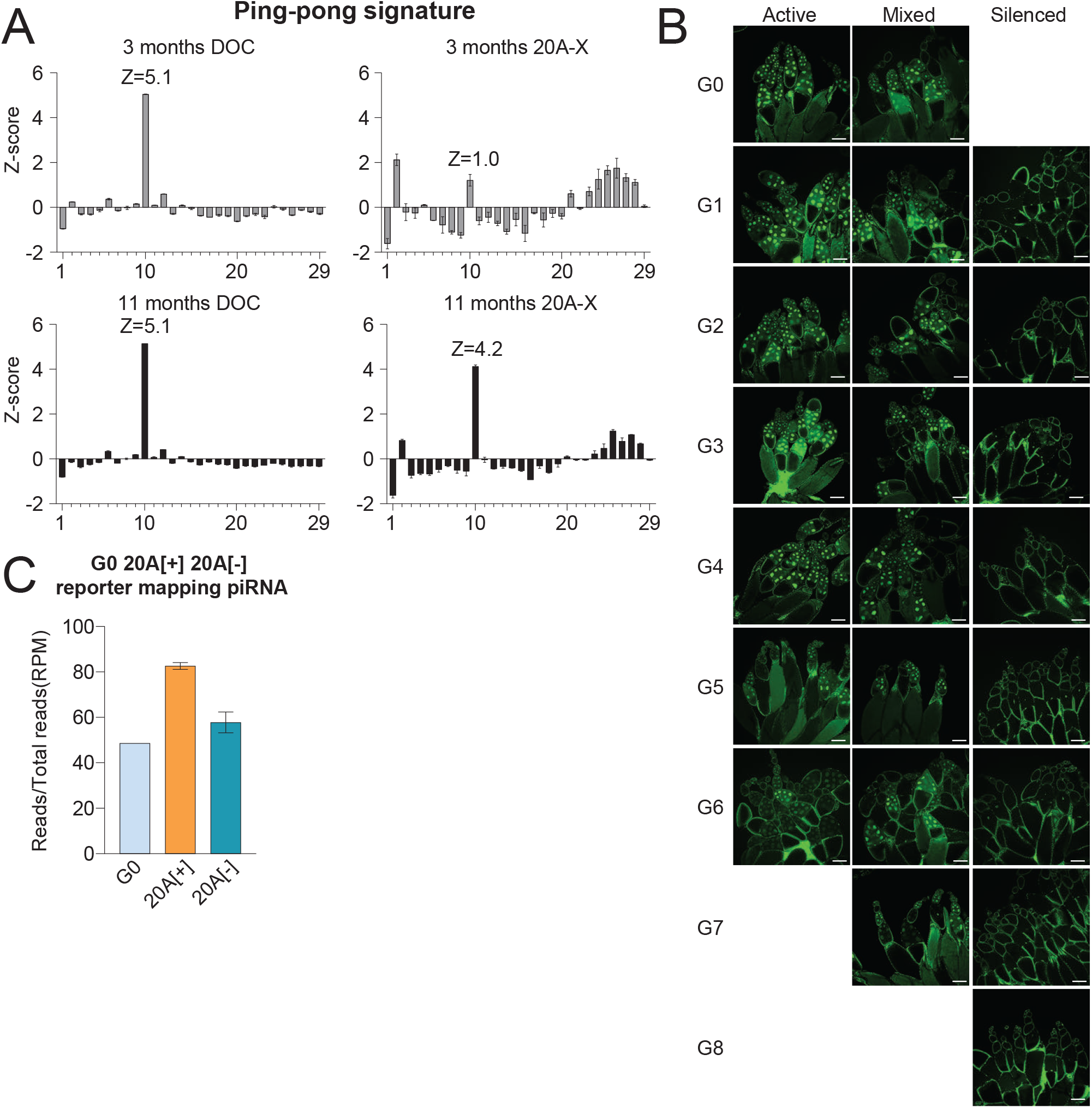
related to Figure 5. **(A)** Analysis of ping-pong signature (overlap between 5’ends of piRNA mapping in opposite orientation) of piRNAs mapping to 20A-X and DOC piRNAs. Error bars indicate standard deviation of two biological replicates. **(B)** GFP expression in ovary of each generation (G0 to G8) from the crosses shown on Fig. 5D. Scale bar is 100µm. **(C)** Comparison of the number of reporter-mapping piRNAs in small RNA libraries from ovaries of 20A-X flies after paternal transmission (G0) (after normalized to copy number based on GFP qPCR results) and in small RNA libraries from ovaries of 20A[+] and 20A[-] flies.

**Supplementary Table 1.** *Drosophila melanogaster* stocks.

| Stock | Source | Identifier |
| --- | --- | --- |
| pUbi > eGFP-NLS 20A[-] | This study | N/A |
| pUbi > eGFP-NLS 20A[+] | This study | N/A |
| pUbi > eGFP-NLS 20A-X | This study | N/A |
| pUbi > eGFP-NLS 42AB[-] | This study | N/A |
| pUbi > eGFP-NLS 66A6 | This study | N/A |
| pUbi > eGFP(sense)-NLS-GFP(antisense) 20A[-] | This study | N/A |
| pUbi > eGFP(sense)-NLS-GFP(antisense) 20A[+] | This study | N/A |
| pUbi > eGFP(sense)-NLS-GFP(antisense) 66A6 | This study | N/A |
| MI07308 | Bloomington stock | BDSC #43121 |
| MI08972 | Bloomington stock | BDSC #50496 |
| UASp > small hairpin white | Bloomington stock | BDSC #33623 |
| UASp > small hairpin Rhino | Bloomington stock | BDSC #35171 |
| UASp > small hairpin Zucchini | VDRC | #313693 |
| 20A-X deletion gRNA | This study | N/A |
| Δ20A-X | This study | N/A |
| 20A promoter gRNA | This study | N/A |
| Δ20A promoter | This study | N/A |
| Δ20A promoter 20A-X | This study | N/A |
| attP40{nos-Cas9} | NIG-FLY | CAS-0001 |
| Maternal alpha-tubulin67C > Gal4 | Bloomington stock | BDSC #7063 |
| pNanos > Gal4 | Bloomington stock | BDSC #4937 |

**Supplementary Table 2.**
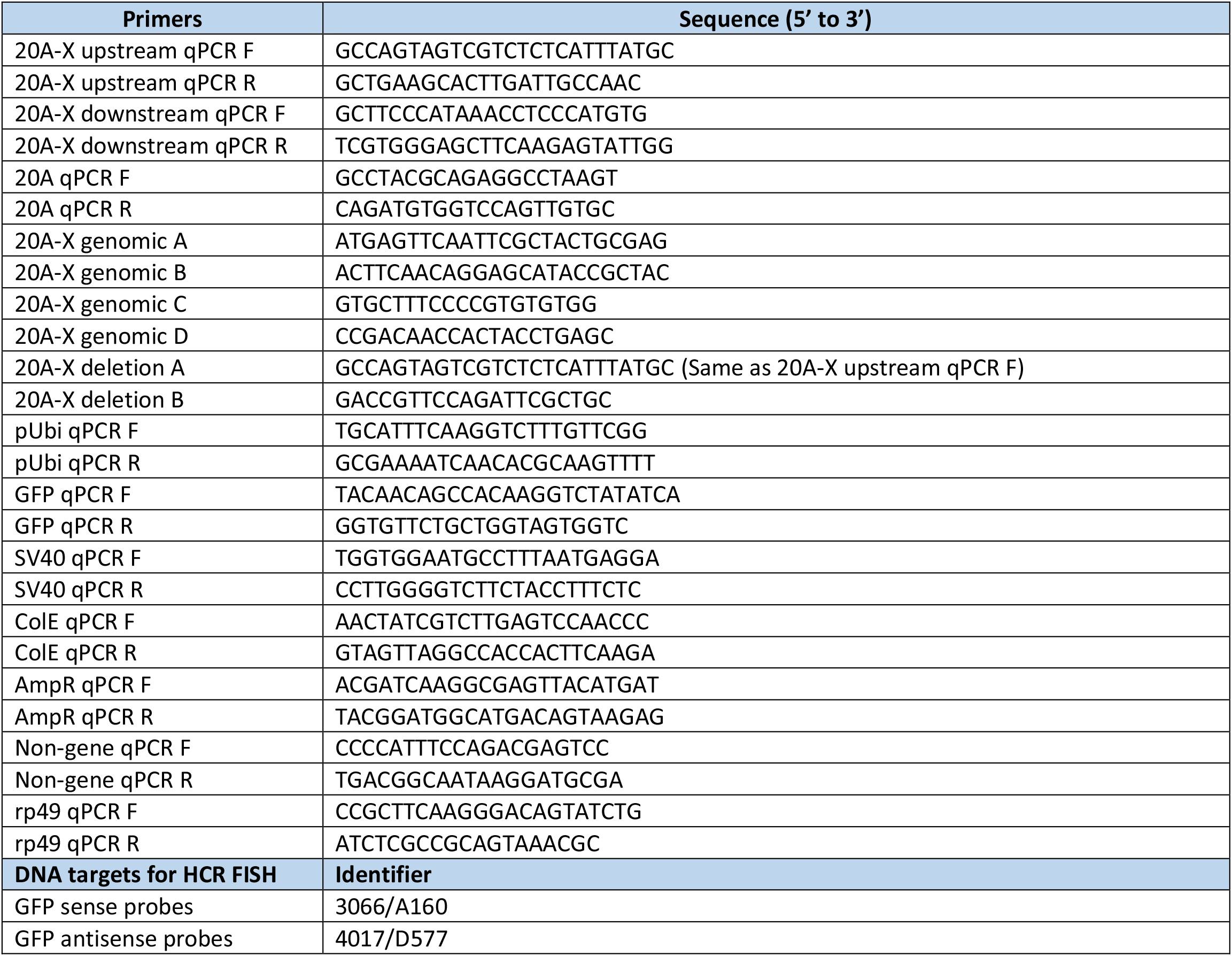
Primers.

## Materials and Methods

### *Drosophila* stocks

All flies were raised at 25°C. The shWhite (BDSC #33623) and shRhino (BDSC #35171) stocks were obtained from Bloomington, shZucchini (#313693) stock was obtained from the Vienna *Drosophila* Resource Center, *Ago2*^*414*^ (#109027) stock was obtained from Kyoto Stock Center. shRNAs were driven by the nos-GAL4 driver (BDSC #4937).

### Transgenic flies

To make the Ubi-GFP-NLS-SV40 reporter, Ubiquitin promoter, GFP-NLS and SV40 were PCR amplified and PCR products were assembled into the EcoR1 and BamH1 digested pBS-KS-attB1-2 vector by Gibson Assembly. The recombinant vector was integrated into three genomic sites chrX: 21522657 dm6 (20A, BDSC #50496), chr2R: 6338399 dm6 (42AB, BDSC #43121) and chr3L: 7575013 dm6 (control, BDSC #38579). To make the Ubi-GFP(sense)-NLS-GFP(antisense)-SV40 fly, antisense GFP was amplified by PCR and digested with BglII and EagI, then ligated into the BglII and EagI double-digested Ubi-senseGFP vector. Recombinant vectors were integrated into genomic site chrX: 21522657 dm6 (20A, BDSC #50496) and chr3L: 7575013 dm6 (control, BDSC #38579). All constructs were injected by Bestgene.

20A-X and 20A promoter deletion gRNAs were designed using CRISPR Optimal Target Finder and synthesized by IDT. Oligos were cloned into the pCFD5 vector by Gibson Assembly as described(Port and Bullock, 2016). The DNA oligos sequences are shown below:

20A-X deletion gRNA sequence Forward:

GCGGCCCGGGTTCGATTCCCGGCCGATGCA<u>TTGAAGCTCCCACGAAGTTA</u>GTTTTAGAGC TAGAAATAGCAAG

Reverse: ATTTTAACTTGCTATTTCTAGCTCTAAAAC<u>TAGTTGACGAGTGTCCGCTT</u>TGCACCAGCCGG GAATCGAACCC

20A promoter gRNA sequence Forward:

GCGGCCCGGGTTCGATTCCCGGCCGATGCA<u>ACTACGTTACTAAGCATTTG</u>GTTTTAGAGCT AGAAATAGCAAG

Reverse: ATTTTAACTTGCTATTTCTAGCTCTAAAAC<u>GATGTCCAAACTTGCAATTT</u>TGCACCAGCCGG GAATCGAACCC

All transgenic constructs were inserted into the attP40 landing site at 25C6 (y1w67c23; P[CaryP]attP40) on the 2^nd^ chromosome and attP2 landing site at 68A4 (y1w67c23; P[CaryP]attP2) on the 3^rd^ chromosome, unless specifically mentioned. To obtain 20A promoter deletion flies, flies carrying gRNAs were crossed with Nos-Cas9 flies (CAS-0001, NIG-FLY). Individual progeny were screened to verify the promoter deletion by genomic PCR followed by sanger sequencing. Transgenic flies used in this study are listed in Supplementary Table 1.

### RNA HCR-FISH

The HCR-FISH RNA protocol was adapted from a previous protocol(Choi et al., 2018; Luo et al., 2020). The probes were designed and synthesized by Molecular Technologies and Alexa594 was used for probe detection. Images were acquired using the ZEISS LSM880 and data was processed using Zen software.

### ChIP-seq and ChIP-qPCR

All ChIP experiments were performed based on previous description(Chen et al., 2016), using anti-H3K9me3 antibody from Abcam (ab8898) and anti-Rhino antibody obtained from the Brennecke lab. SYBR Green qPCR was performed by using MyTaq HS Mix (BioLine). CT values were calculated from technical duplicates. All ChIP-qPCR were normalized to respective inputs and to control region rp49. ChIP-qPCR were performed on a Mastercycler^®^ep Realplex PCR thermal cycler machine (Eppendorf), All qPCR primers are listed in Supplementary Table 2. ChIP-seq libraries were generated using the NEBNext ChIP-Seq Library Prep Master Mix Set. All libraries were sequenced on the Illumina HiSeq 2500 platform (SE 100 bp reads).

### Small RNA-seq

Total RNA was isolated from dissected ovaries using TRIzol (ThermoFisher #15596018). 4µg total RNA was loaded onto a 15% polyacrylamide gel and small RNA between 19 and 29 nt in length was excised and isolated. Size selected small RNA was ethanol-precipitated and small RNA library constructed using the NEBNext small RNA library preparation set (#E7330S). Libraries were sequenced on the Illumina HiSeq 2500 platform (SE 50-bp reads).

### RT-qPCR

Around 20 ovaries were dissected and homogenized in 1mL TRIzol (ThermoFisher #15596018) and total RNA was extracted following the manufacturer’s recommendation. DNAase I treatment and reverse transcription was performed from 1µg total RNA starting material, using DNase I and SuperScript III (Invitrogen) following the manufacturer’s recommendation. qPCR was performed by using MyTaq HS Mix (BioLine) contain SYBR Green on a Mastercycler^®^ep Realplex PCR thermal cycler machine (Eppendorf). CT values were calculated from technical duplicates. All qPCR data were normalized to the *rp49* mRNA expression. All qPCR primers are listed in Supplementary Table 2.

### DNA FISH

Polytene chromosomes DNA FISH was performed as previously described(Cai et al., 2010; Lavrov et al., 2004) with the following modifications. Salivary glands were dissected from 3^rd^ instar larvae and fixed in fixation buffer (3.7% Formaldehyde, 1% Triton X-100 in PBS, pH 7.5) for 5 min, then transferred into solution (3.7% Formaldehyde, 50% acetic acid) for 2 min on the cover slip. Cover slip was put on poly-L-lysine coated microscope slide and chromosomes were spread and quashed by gently moving the cover slip back and forth followed by pressure applied to the cover slip by thumb. Slides were flash frozen in liquid nitrogen to remove cover slip and submerged in PBS for 10 min followed by three 5 min washes in 2x SSC. Samples were dehydrated by 5 min incubations twice in 70% ethanol and twice in 96% ethanol, followed by air-drying slides. Slides were incubated in 2x SSC for 45 min at 70°C, and dehydrated again as described above. To denature the DNA, slides were incubated in 100 mM NaOH for 10 min, washed three times with 2x SSC and dehydrate as described above. Slides were incubated in hybridization buffer (2X SSC, 10% dextran sulfate, 50% formamide, 0.8 mg/mL salmon sperm DNA) for 5 min at 80°C and snap cooled on ice. DNA FISH probes were prepared following the manufacturer’s recommendations (ThermoFisher # F32947 and F32949) using the BAC construct (BACPAC Resources #CH322-184J4) as probe template for 20A (Alexa 594) and the original reporter vector (non-RMCE) as probe template for the GFP reporter (Alexa 488). Probes pre-warmed to 37°C were loaded on the slides, covered with cover slip, sealed with rubber cement and incubated in a dark and humid chamber at 37°C overnight. Slides were washed in 2x SSC three times at 42°C and once at RT, 5 min each time followed by DAPI staining for 10 min and two washes in PBS. Slides were mounted with mounting medium (Vector Labs #H-1000). Images were acquired using the ZEISS LSM880.

### Bioinformatic analysis

ChIP-seq processing and mapping. Trimmomatic (version 0.33)(Bolger et al., 2014) and cutadapt (version 1.15)(Martin, 2011) were used to trim off adaptors and filter out those shorter than 50 nt after trimming. The first 50 nt from each read were mapped to dm3 genome and vector sequence respectively, using Bowtie(Langmead et al., 2009) (version 1.0.1, parameters: -v 2 -k 1 -m 1 -t --best -y --strata). After mitochondria reads were removed, aligned reads were then used to generate piled-up RPM signals and enrichment profiles by our customized scripts and deepTools(Ramirez et al., 2016). Regions blacklisted by ENCODE(Amemiya et al., 2019) were excluded from enrichment analysis. Read counts over equal-sized bins were calculated using deepTools2 and BEDOPS(Neph et al., 2012), and figures were made using Matlab. All the scripts we used can be found on GitHub (https://github.com/brianpenghe/Luo_2021_piRNA/blob/main/ChIP-seq.md).

To map fusion read, the first 20 nt and full length of the reads were mapped to vector sequences using the aforementioned bowtie settings. Reads where the first 20nt mapped to vector sequences but the full length did not were selected. The last 20 nt of such reads was mapped to the reference genome with the same settings. Mappable reads among these were considered fusion reads between the vector and genome which were used to identify insertion location. The scripts are available on GitHub (https://github.com/brianpenghe/Luo_2021_piRNA/blob/main/FusionReads.md).

For small RNA-seq analysis, Trimmomatic and cutadapt were used to trim off adaptors and filter out reads shorter than 20 nt after trimming. We then extracted reads of have specific lengths were extracted: 21-22 nt (siRNA), 23-29 nt (piRNA) and 21-30 nt (small RNA). The selected reads were mapped to the dm6 genome using Bowtie (parameters: -v 0 -a -m 1 -t --best --strata). After mitochondrial reads were removal, deepTools2 and BEDOPS were used to calculate read counts over equal-sized bins. Ping-pong signature was inferred using a published method(Antoniewski, 2014). Reporter coverage was calculated based on 10 nt bin size, and figures were made using Matlab. The mapping scripts are available on GitHub (https://github.com/brianpenghe/Luo_2021_piRNA/blob/main/piRNA-seq.md).

## Data availability

Libraries generated from this study are deposited in GEO under accession codes GSE193091. The scripts are available on GitHub: https://github.com/brianpenghe/Luo_2021_piRNA. Pol ll ChIP-seq data analyzed in this study were from GSE43829 (Le Thomas et al., 2013) and GSE97719 (Andersen et al., 2017).

## Author Contributions

Y.L., K.F.T. and A.A.A. designed experiments. Y.L. performed all experiments; N.K. helped to perform imaging and ChIP experiments; P.H. performed the computational analysis; Y.L. and P.H. analyzed the data; Y.L., K.F.T. and A.A.A. wrote the paper.

